# Identification and targeting of a unique NaV1.7 domain driving chronic pain

**DOI:** 10.1101/2022.07.09.499431

**Authors:** Kimberly Gomez, Harrison J. Stratton, Paz Duran, Santiago Loya, Cheng Tang, Aida Calderon-Rivera, Liberty François-Moutal, May Khanna, Cynthia L. Madura, Shizhen Luo, Dongzhi Ran, Lisa Boinon, Samantha Perez-Miller, Aubin Moutal, Rajesh Khanna

## Abstract

Despite identification of several small molecules directly targeting the voltage-gated sodium channel NaV1.7, none has been clinically successful. We reported that preventing addition of a small ubiquitin-like modifier (SUMO) on the NaV1.7-interacting cytosolic collapsin response mediator protein 2 (CRMP2) blocked NaV1.7 functions and was antinociceptive in rodents. Here, we discovered a 15 amino acid CRMP2 regulatory sequence (CRS) unique to NaV1.7 that is essential for this regulatory coupling. CRMP2 preferentially bound to the NaV1.7 CRS over other isoforms. Substitution of the NaV1.7 CRS with the homologous domains from the other eight voltage-gated sodium channel isoforms decreased tetrodotoxin-sensitive NaV1.7 currents in rodent sensory neurons. A cell-penetrant version of NaV1.7-CRS reduced NaV1.7 currents and trafficking, decreased presynaptic NaV1.7 localization, reduced spinal neurotransmitter release, and reversed mechanical allodynia in a rat spared nerve injury model of neuropathic pain. Interfering with NaV1.7-CRMP2 coupling did not produce motor impairment and spared thermal, inflammatory, and post-surgical nociception. As proof-of-concept for NaV1.7-targeted gene therapy, we found that NaV1.7-CRS packaged into an adeno-associated virus recapitulated the effects on NaV1.7 function in both rodent and rhesus macaque sensory neurons and both reversed and prevented the development of mechanical allodynia in a neuropathic pain model in male and female rodents.

**One Sentence Summary:** A novel regulatory domain on the voltage gated sodium channel NaV1.7 that can be targeted to produce analgesia.

## INTRODUCTION

The generation of action potentials relies in part on the threshold set by voltage gated sodium channels such as NaV1.7. Encoded by the SCN9A gene, NaV1.7 is expressed in nociceptive sensory neurons of the dorsal root ganglion (DRG) (*1*) and was established as a major contributor to human pain signaling (*1, 2*). Patients carrying activating mutations on SCN9A are burdened with painful heritable syndromes such as inherited erythromelalgia, small-fiber neuropathy, and paroxysmal extreme pain disorder (*3*). Conversely, *SCN9A* loss-of-function mutations produce congenital insensitivity to pain (*4*). NaV1.7 function is increased in pre-clinical rodent models of chronic neuropathic pain (*5*)(*6*)(*7*). While this evidence points to a critical role for NaV1.7 in pain, direct inhibition of this channel failed to reach satisfactory clinical endpoints in clinical trials (*1, 8–14*).

We endeavored to understand how NaV1.7 is trafficked and retained at the pre-synapse in primary afferents. We identified the collapsin response mediator protein 2 (CRMP2) as a critical protein regulating NaV1.7 membrane localization (*15–18*). When phosphorylated by cyclin dependent kinase 5 (Cdk5) and SUMOylated, CRMP2 protected NaV1.7 from internalization to maintain its pre-synaptic localization in chronic neuropathic pain (*19, 20*). Pharmacological inhibition of CRMP2 phosphorylation or SUMOylation, specifically decreased NaV1.7 membrane localization and function to provide pain relief in rodent models of chronic neuropathic pain (*20, 21*). However, the mechanism driving the selective regulation of NaV1.7 by CRMP2 was not elucidated.

Using peptide mapping, we identified a unique CRMP2 binding domain in the first intracellular loop of NaV1.7. We further demonstrate that this CRMP2 regulatory sequence (CRS) is necessary for NaV1.7 membrane localization and function. Targeting CRS with a decoy peptide fused to a cell penetrating sequence or genetically encoded, inhibited NaV1.7 in vitro and in vivo to block chronic neuropathic pain, but not physiological pain. Finally, we validated the translational potential of the CRS domain on NaV1.7 in sensory neurons and spinal cord from non-human primates. Together, our results reveal a unique domain (CRS) amenable to therapeutic targeting to specifically inhibit NaV1.7.

## RESULTS

### Identification of a unique NaV1.7 regulatory domain

The NaV1.7 intracellular domains mediating channel regulation are not yet well understood (*22–24*). To identify potential sites of interaction between CRMP2 and NaV1.7, we generated a peptide array spanning the intracellular loops of human NaV1.7 (**Fig. 1A**). We hybridized the array with spinal cord lysate from rat with chronic neuropathic pain (spared nerve injury, SNI) and found that CRMP2 bound to peptide number 141 (**Fig. 1B**) (#141; hNaV1.7, 706-SRGKCPPWWYRFAHK-720). CRMP2 bound the same NaV1.7 peptide in lysates from porcine and human DRG or spinal cord (**Fig. 1B**). Because CRMP2 regulates NaV1.7 exclusively (*19, 21*), we tested if CRMP2 could bind analogous hNaV1.x-CRS regions (**Fig. 1C**). CRMP2 bound to hNaV1.7-CRS with an affinity six-fold higher than to any other hNaV1.x peptides: KD=0.99 μM for hNaV1.7 versus K_D_=6.18 μM for hNaV1.3 and K_D_=6.28 μM for hNaV1.4 (Fig. 1C, Fig. S1). No binding was detected to other hNav1.x-CRS homologous domains (**Fig. 1C**, **Fig. S1**).

**Figure 1.**
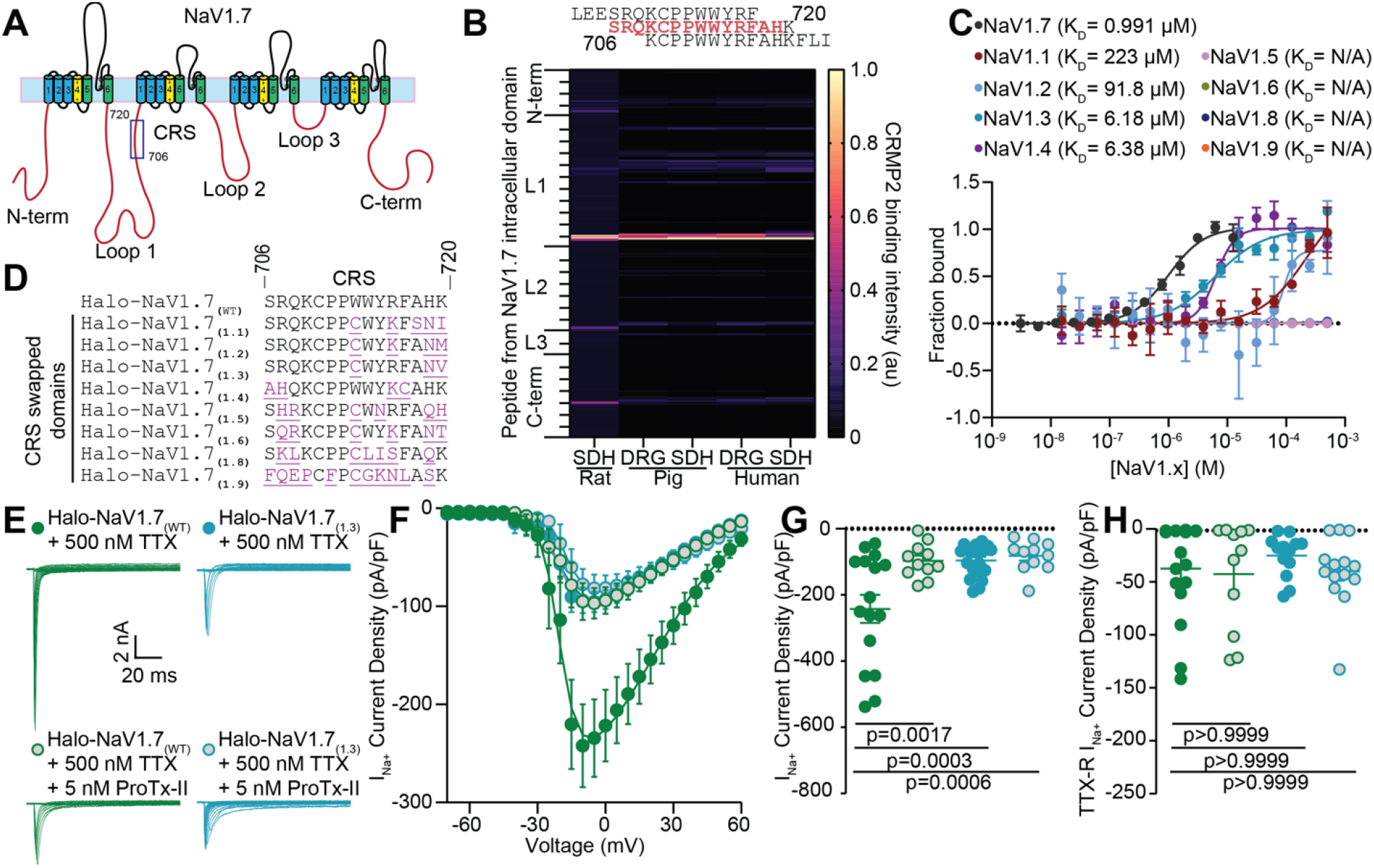
Identification of a unique CRMP2 regulatory domain on NaV1.7. (**A**) Cartoon of the domain structure of human NaV1.7 with intracellular loops labeled. These loops were divided into 384 15-mer peptides with 12 overlapping amino acids and were printed onto a peptide array. (**B**) Fluorescent intensity of CRMP2 binding to the peptide array from rat, pig and human DRG and spinal cord lysate (n=4). (**C**) Microscale thermophoresis of NTA-labeled His-CRMP2 (200 nM) with different concentrations of the CRMP2-binding peptide from NaV1.7 or the homologous region of NaV1.X channels, fitted to a one-site binding model (r^2^= 0.97 for NaV1.7 and 0.76 for NaV1.1; n = 4; Bottom). (**D**) Sequence alignment of the CRMP2 binding domain on NaV1.7 with the CRS analogous domains. (**E**) Representative traces of sodium currents recorded from DRGs transfected with Halo-NaV1.7_(WT)_ (green) and Halo-NaV1.7_(1.3)_ (teal) in the presence or absence of 5 nM ProTx-II. (**F**) Current density-voltage relationship for Halo-NaV1.7_(WT)_, Halo-NaV1.7_(1.3)_, and both groups following treatment with 5 nM ProTx-II. (**G**) Peak Halo-NaV1.7 current density for Halo-NaV1.7_(WT)_, Halo-NaV1.7_(WT)_ + 5 nM ProTx-II, Halo-NaV1.7_(1.3)_, and Halo-NaV1.7_(1.3)_ + 5 nM ProTx-II. (**H**) TTX-R peak current density for the conditions listed above. Each group was compared to its own on-plate Halo-NaV1.7_(WT)_ control group and statistical significance was determined using a Mann-Whitney test. n=11-16 cells; error bars indicate mean ± SEM; *p* values as indicated (**Table S5**); Kruskal-Wallis test with the Dunn post hoc test. All biophysical parameters are shown in **Table S1**.

To test if NaV1.7-CRS is responsible for the selective regulation of NaV1.7, we used a plasmid encoding a mNaV1.7 channel with resistance to tetrodotoxin (Y362S, TTX-R) (*25*) and added a N-terminal extracellular HaloTag reporter linked to NaV1.7 via the transmembrane helix from the β4 subunit of VGSC (referred here as Halo-NaV1.7_(WT)_) (*26*). We mutated the CRS domain in Halo-NaV1.7_(WT)_ to the homologous CRS sequences from other Nav1.x channels (**Fig. 1D**). All NaV1.7 mutants expressed well in mouse catecholamine A differentiated (CAD) cells (**Fig. S2A**) and could be detected with Halo ligand fluorescence in transfected rat DRG neurons (**Fig. S2B**). We recorded sodium currents for each Halo-NaV1.7_(WT/1.x)_ construct expressed in rat DRG neurons where the endogenous TTX-sensitive currents were blocked (500nM TTX) to isolate solely the currents arising from transfected channels (**Fig. 1E-G, Fig. S3**). We used changes in sodium current density to infer the degree of regulation imposed by substitution of the NaV1.7-CRS. We found that all of our mutations in the CRS domain on Halo-NaV1.7 imposed a profound reduction of current density (**Fig. S3**). To test if all the Halo-NaV1.7 current was nonfunctional when the CRS was mutated, we applied the NaV1.7 selective blocker ProTx-II which did not result in further reduction of current density (**Fig. 1E-G**). This experiment was done with Halo-NaV1.7_(1.3)_ compared to Halo-NaV1.7_(WT)_ to test this idea using the CRS domain with the highest homology with the CRS_(1.7)_ sequence. Importantly, the current density of TTX-R channels - NaV1.8 and NaV1.9 - was not affected in any of these experiments (**Fig. 1H**, **Fig. S3**). The activation and inactivation constants of NaV1.7 remained unchanged in our experiments (**Table S1**, **Fig. S3**). These results identify a unique regulatory site necessary for NaV1.7 function in DRG sensory neurons.

### Competitive inhibition of CRMP2 binding to NaV1.7 decreases Na^+^ current

Having discovered a novel regulatory domain on NaV1.7, we next tested if this sequence could be targeted to limit NaV1.7 function and mitigate pain in vivo. To this end, a peptide corresponding to the NaV1.7-CRS fused to the cell penetrating trans-activator of transcription (TAT) sequence (YGRKKRRQRRR) (*27*) was synthesized. To increase the juxtamembrane concentration of this decoy peptide, we added a 14-carbon myristate group (Myr) to the N-terminus as an anchor to the plasma membrane (*28*) (Fig. 2A). At 5μM, the Myr-TAT-NaV1.7-CRS peptide could disrupt CRMP2 interaction to the CRS domain forming peptides on our peptide array (**Fig. S4**). Myr-TAT-NaV1.7-CRS was also able to inhibit the CRMP2/NaV1.7 interaction in CAD cells (**Fig. 2B-C**). To ensure that the actions of Myr-TAT-NaV1.7-CRS is not via an effect on CRMP2 post-translational modifications known to affect its interaction with NaV1.7 (*19, 29*), we verified that Myr-TAT-NaV1.7-CRS had no effect on CRMP2 SUMOylation, phosphorylation by Cdk5 (pS522) and by GSK3β (pT514) (**Fig. S5**). In DRG neurons, acute application of 5 μM Myr-TAT-NaV1.7-CRS reduced sodium current density (**Fig. 2D-F**) without affecting activation or steady-state inactivation kinetics (**Table S2**) or TTX-R current density, compared to Myr-TAT-SCR (**Fig. 2G**). To test if all of the current carried by NaV1.7 is inhibited following treatment with Myr-TAT-NaV1.7-CRS, we added ProTx-II (5nM) to block any remaining NaV1.7 channels. While ProTx-II decreased sodium current density in DRG neurons treated with the scramble peptide control, co-application with Myr-TAT-NaV1.7-CRS did not produce any further current reduction (**Fig. 2H-J**), thus, demonstrating that Myr-TAT-NaV1.7-CRS achieved maximal NaV1.7 blockade in DRG neurons. We next used our validated siRNA (*19*) to silence CRMP2 expression in DRG neurons and test if Myr-TAT-NaV1.7-CRS could reduce NaV1.7 currents independently of CRMP2. As before, reducing CRMP2 expression inhibited peak sodium current density by ~43% (**Fig. 2K-M**). Co-treatment with Myr-TAT-NaV1.7-CRS failed to elicit a further reduction of current density (**Fig. 2K-M**). Furthermore, we found that Myr-TAT-NaV1.7-CRS had no effect on voltage-gated potassium (**Fig. S6A-C**) or calcium channels (**Fig. S6D-G**) in DRGs. The above results demonstrate that the CRS domain can be functionalized to specifically inhibit NaV1.7 in DRG sensory neurons.

**Figure 2.**
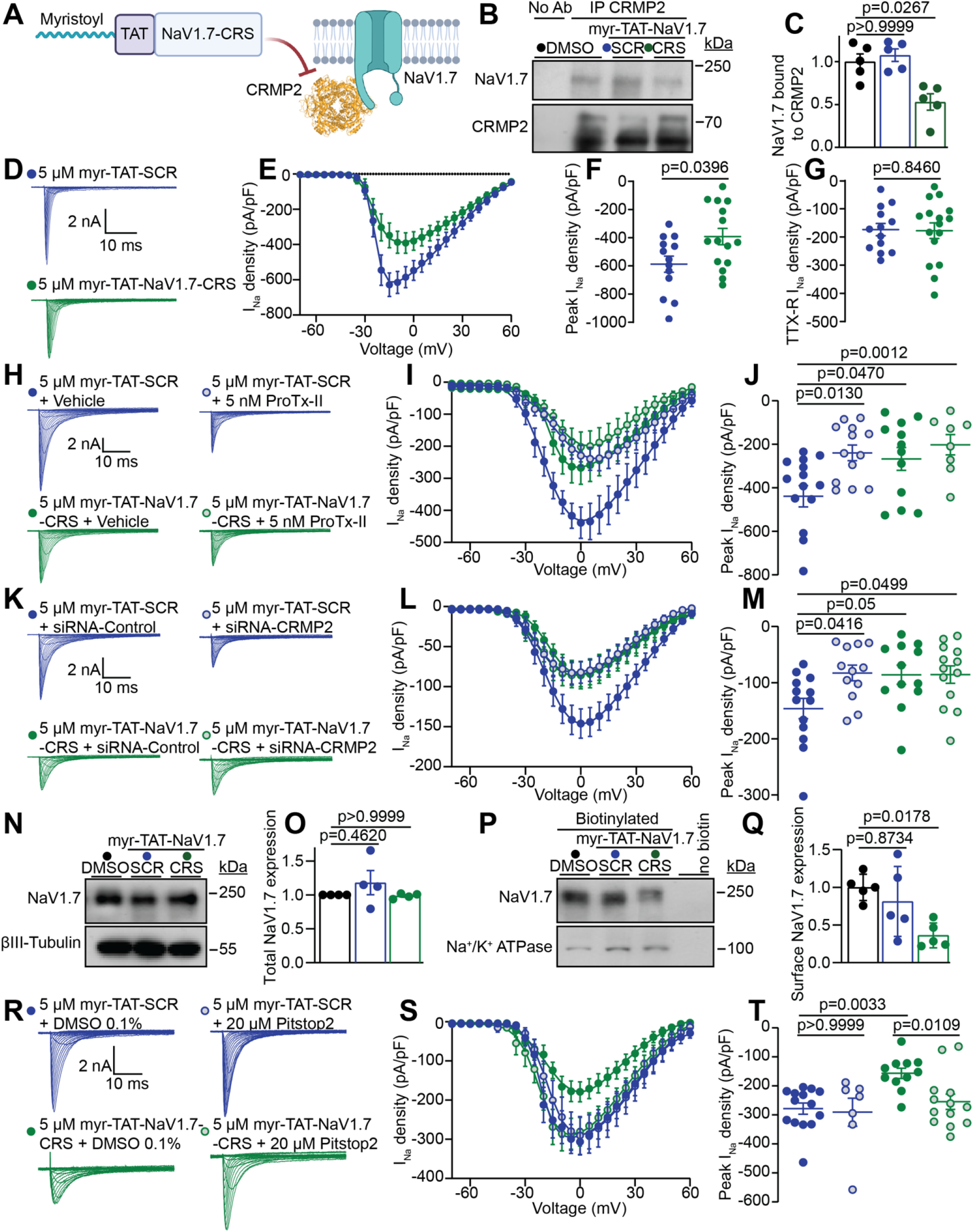
Myr-TAT-NaV1.7-CRS causes CRMP2 dependent reduction in NaV1.7 currents in DRG neurons. (**A**) The Myr-TAT-NaV1.7-CRS peptide competes for binding to NaV1.7 regulatory proteins. (**B**) Representative immunoblots and summary (**C**) of CRMP2 immunoprecipitation (IP) to detect NaV1.7 from CAD cells treated with the indicated peptides (n= 5). (**D**) Representative current traces recorded from small-sized DRG neurons in the presence of 5 μM Myr-TAT-SCR peptide (blue circles, n=15) and Myr-TAT-NaV1.7-CRS peptide (green circles, n=16). (**E**) Summary of Boltzmann fits for current density-voltage curves, (**F**) peak current densities and (**G**) electrically isolated TTX-R currents. (**H**) Representative current traces recorded from small-sized DRGs incubated with Myr-TAT-SCR and Myr-TAT-NaV1.7-CRS in the presence and absence of the NaV1.7 specific inhibitor ProTx-II. (**I**) Boltzmann fits of the current density-voltage curves for each treatment group. (**J**) Summary of peak current densities. N=8-12 cells; (**K**) Representative current traces recorded from small-diameter DRGs transfected with either siRNA-Control or siRNA-CRMP2 and treated with Myr-TAT-NaV1.7-CRS or Myr-TAT-SCr as indicated. (**L**) Boltzmann fits for current density-voltage curves and (**M**) Peak current densities. n=12-13 cells; (**N**) Representative immunoblots and (**O**) summary (bottom) of NaV1.7 expression in CAD cells treated with the indicated peptides. ßIII-Tubulin is used as a loading control (n = 4). (**P**) Representative immunoblots of streptavidin-enriched surface fractions probed for NaV1.7 and Na+/K+ ATPase as a control. (**Q**) bar graph with scatter plot of mean surface localized NaV1.7 in CAD cells treated with the indicated peptides. (**R**) Representative current traces recorded from small diameter DRG neurons in the presence of Myr-TAT-SCR or Myr-TAT-NaV1.7-CRS ± 20 μM Pitstop2 as indicated; (**S**) Boltzmann fits for current density-voltage curves and (**T**) Summary of peak current densities (pA/pF) showing pitsopt2 blocked the current reduction imposed by Myr-TAT-NaV1.7-CRS. n=7-14 cells; error bars indicate mean ± SEM;*p* values as indicated; Kruskal–Wallis test with Dunnett’s post hoc comparisons (**Table S5**). All biophysical parameters are shown in **Table S2**.

### Disruption of the CRS domain induces NaV1.7 internalization

CRMP2 binding to NaV1.7 maintains the membrane localization of the channel (*29*), so we next examined if blocking the CRS domain on NaV1.7 could induce its internalization. Myr-TAT-NaV1.7-CRS did not change NaV1.7 protein expression (**Fig. 2N-O**). In cell surface biotinylation experiments, Myr-TAT-NaV1.7-CRS diminished NaV1.7 surface expression by ~64% (**Fig. 2P-Q**). In DRG neurons, Pitstop2 (20 μM) — an inhibitor of clathrin assembly — rescued the decrease of sodium currents imposed by Myr-TAT-NaV1.7-CRS (**Fig. 2R-T**). No changes in the biophysical properties of activation or inactivation were seen between these four conditions (**Table S2**). These findings show that the reduction of NaV1.7 currents due to the blockade of the CRS domain is elicited via an active internalization of the channel.

### Inhibition of the NaV1.7 CRS domain reduces DRG excitability and spinal CGRP release

NaV1.7 channels set the threshold for action potential firing in sensory neurons (*30*). Therefore, inhibition of NaV1.7 currents by Myr-TAT-NaV1.7-CRS may in turn decrease action potential firing in DRG neurons. Compared to neurons treated with the Myr-TAT-SCR control, neurons treated with Myr-TAT-NaV1.7-CRS had decreased excitability (**Fig. 3A-B**). Their resting membrane potential was not affected (**Fig. 3C**) but the rheobase — the minimum current necessary to evoke a single AP — was increased in DRG neurons treated with Myr-TAT-NaV1.7-CRS (**Fig. 3D-E**). Analysis of the action potential waveform revealed no alterations of peak/antipeak amplitude, rise/decay tau while the time to peak was increased for some current steps (**Fig. S7**). These findings show that targeting the CRS domain can attenuate the intrinsic excitability of DRG neurons.

**Figure 3.**
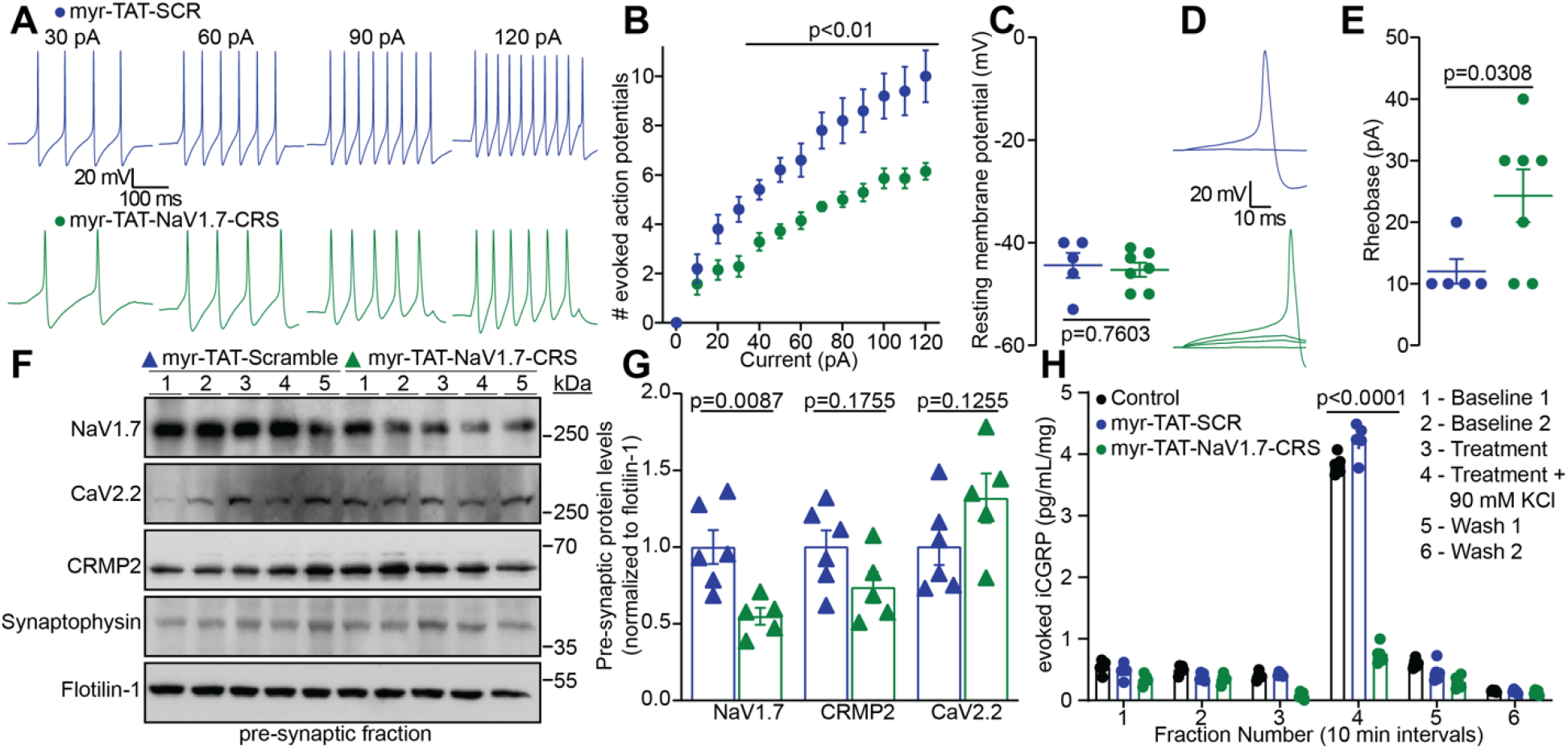
Disruption of the NaV1.7-CRMP2 interaction decreases presynaptic NaV1.7, sensory neuron excitability and spinal cord neurotransmitter release. (**A**) Representative recordings of evoked action potentials recorded from small-diameter DRGs in response to depolarizing current injections of 30, 60, 90 and 120 pA after addition of 5μM Myr-TAT-SCR (blue) and Myr-TAT-NaV1.7-CRS peptides (green). (**B**) Quantification of the number current-evoked action potentials in response to 0-120 pA of injected current. (**C**) Resting membrane potential of cells recorded in A. (**D**) Representative traces and (**E**) quantification showing an increased rheobase in cells treated with Myr-TAT-NaV1.7-CRS. n=5-7 cells; (**F**) Representative immunoblots of NaV1.7, CRMP2 and CaV2.2 expression from the presynaptic fraction of male rat spinal dorsal horn after i.t. injection of 20 μg/5 μl of Myr-TAT-SCR (n=6) or Myr-TAT-NaV1.7-CRS (n=5) peptides. Tissues were collected 1 hour after injection. (**G**) bar graph with scatter plot showing quantification of the data in **F**. Myr-TAT-NaV1.7-CRS specifically decreased NaV1.7 spinal pre-synaptic localization. (**H**) KCl depolarization-evoked CGRP release was measured from spinal cord isolated from naïve male rats following pre- and co-incubation with 0.1% DMSO (control), 5 μM of Myr-TAT-SCR, or 5 μM Myr-TAT-NaV1.7-CRS peptides. Histograms showing immunoreactive CGRP levels observed in bath solution normalized to tissue mass (n=4 animals); error bars indicate mean ± SEM; data was analyzed by either Mann-Whitney test or two-way ANOVA (details are in **Table S5**), *p* values as indicated.

To parallel our in vitro data, we used an in vivo synaptic fractionation approach to assess if targeting the CRS domain of NaV1.7 could also reduce NaV1.7 spinal presynaptic localization. Male rats with SNI were injected with Myr-TAT-NaV1.7-CRS (20 μg/5 μl i.t.) or control, 1 h after intrathecal administration, we collected the lumbar dorsal horn and extracted the pre-synaptic fractions (*20*). We found that blocking the CRS domain of NaV1.7 with Myr-TAT-NaV1.7-CRS reduced NaV1.7 presynaptic localization without impacting CRMP2 or CaV2.2 pre-synaptic levels (**Fig. 3F-G**). Next, we examined if Myr-TAT-NaV1.7-CRS induced decrease of pre-synaptic NaV1.7 would impair the evoked release of the nociceptive neurotransmitter calcitonin gene related peptide (CGRP) (*31*). In an ex vivo assay, lumbar spinal cord was stimulated with a high KCl-induced depolarization and fractions collected for evoked CGRP analysis. We found an ~ 80% decrease of evoked CGRP from spinal cords treated with Myr-TAT-NaV1.7-CRS (5 μM) compared to Myr-TAT-SCR or 0.1% DMSO controls (**Fig. 3H**). Together, these data show that interfering with NaV1.7 CRS domain reduces DRG excitability, spinal pre-synaptic localization of NaV1.7 and consequently spinal CGRP release.

### Disruption of NaV1.7-CRMP2 binding confers antinociception and spares physiological pain without affecting motor control

Thus far, our results uncover a new intracellular domain on NaV1.7 that can be blocked to limit the functions of this channel. Using protein extract from the ipsi- and contra-lateral sides from rats with SNI, we found that the interaction of CRMP2 with the target peptide #141 could be specific of a chronic neuropathic pain state (**Fig. 4A**). We next performed an appraisal of nociceptive behaviors that could be mitigated by our blocking peptide Myr-TAT-NaV1.7-CRS (20 μg/5 μl i.t.). We found no effect on behavioral measures of acute and chemical irritant induced pain (**Fig. S8A**), post-operative (**Fig. S8B**), visceral (**Fig. S8C**) and inflammatory (**Fig. S8D**) pain. Next, we administered Myr-TAT-NaV1.7-CRS (20 μg/5 μl i.t.) to male and female rats with SNI and observed a reversal of the mechanical hypersensitivity induced by the nerve injury (**Fig. 4B-C**). NaV1.7 loss in mice leads to a profound loss of pain sensation and elevated nociceptive thresholds (*32, 33*). We found that the intrathecal injection of Myr-TAT-NaV1.7-CRS in male and female mice had no effect in the hot plate (**Fig. 4D**) and tail-flick assays (**Fig. 4E**). Finally, we show that Myr-TAT-NaV1.7-CRS does not impact motor functions, a common unwanted effect of painkillers, measured in the rotarod test (**Fig. 4F**). Altogether, our results demonstrate that the CRS domain can be targeted to provide pain relief in chronic pain conditions without altering physiological pain sensations.

**Figure 4.**
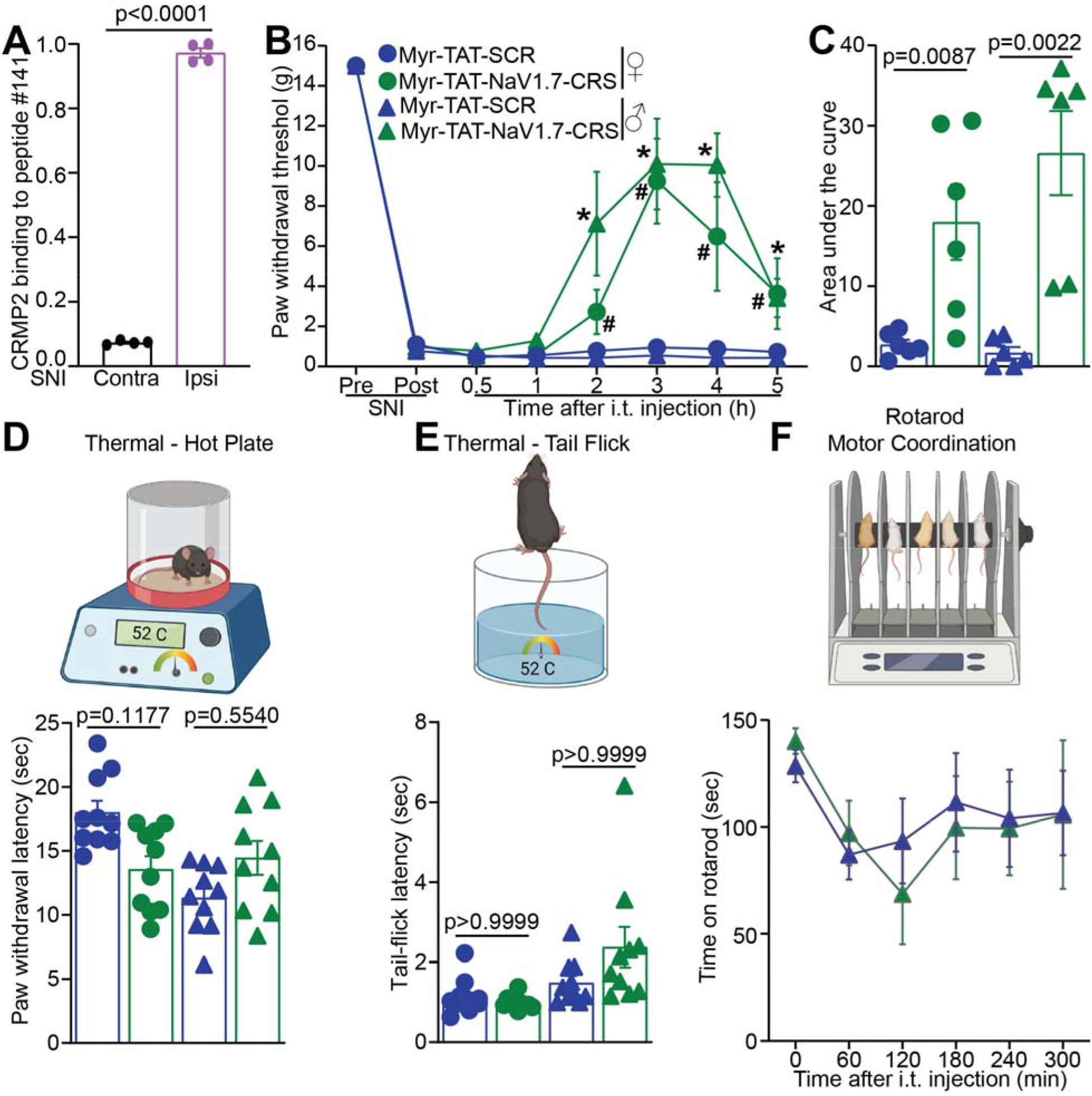
Disruption of the NaV1.7-CRMP2 interaction alleviates mechanical allodynia without affecting physiological pain. (**A**) CRMP2-binding intensity to NaV1.7-derived peptide #141 from contralateral (Contra) and ipsilateral (Ipsi) spinal cords of male rats taken two weeks following SNI (n= 4). (**B**) Time course for male and female rats following administration of Myr-TAT-SCR, or Myr-TAT-NaV1.7-CRS (20μg in 5μl, i.t.). (**C**) Area under the curve for paw withdrawal thresholds showing that Myr-TAT-NaV1.7-CRS reversed mechanical allodynia, n=6 rats; (**D**) Cartoon and bar graph with scatter plot of paw withdrawal latency in the hot plate test (52°C) (n=10 mice). (**E**) Cartoon and bar graph with scatter plot of the tail flick (52°C) test showing no effect of the Myr-TAT-NaV1.7-CRS (n=10 mice). (**F**) Cartoon of the rotarod apparatus to assess motor coordination in rodents. Bar graph with scatter plot showing the latency to fall off a rotating rod was not different between the treatments. n=7 rats; error bars indicate mean ± SEM; Mann-Whitney, Kruskal-Wallis or two-way ANOVA (details are in **Table S5**), *p* values as indicated. The experiments were conducted by investigators blinded to treatments.

### AAV plasmid encoding NaV1.7-CRS reduces NaV1.7 currents, sEPSC frequency and amplitude in DRG neurons

Having demonstrated the therapeutic potential of the CRS sequence in control of NaV1.7 function in vivo, we next asked if a gene therapy approach could result in long term analgesic effects. In an adeno associated virus (AAV) vector, we inserted the CRS sequence at the N-terminus of a GFP (*34*) (AAV-NaV1.7-CRS) to allow tracking of transduced cells. To validate that our construct is functional, we first transfected DRG neurons and recorded sodium currents. DRG neurons expressing AAV-NaV1.7-CRS showed a ~70% decrease in peak sodium current density compared to neurons transfected with the scrambled control plasmid (AAV-SCR) (**Fig. 5A-C**). The fraction of current carried by TTX-R channels (**Fig. 5D**) and the biophysical properties of steady-state inactivation and voltage-dependent activation were not affected by AAV-NaV1.7-CRS (**Table S3**). We next used ProTx-II to test if the reduction imposed by AAV-NaV1.7-CRS was due to NaV1.7. Indeed, ProTx-II did not further block sodium current in AAV-NaV1.7-CRS transfected cells (**Fig. 5E-G**). Finally, we used Pitstop2 to test if the inhibition of NaV1.7 was elicited through the same mechanism as for the Myr-TAT-NaV1.7-CRS peptide (**Fig. 2R-T**). In DRG neurons transfected with AAV-NaV1.7-CRS, Pitstop2 rescued the inhibition of NaV1.7 current compared to the AAV-SCR control (**Fig. 5H-J**). These results validate that the genetic expression of the CRS domain recapitulates the inhibition of NaV1.7 in DRG neurons that we observed with the Myr-TAT-NaV1.7-CRS peptide (**Fig. 2**).

**Figure 5.**
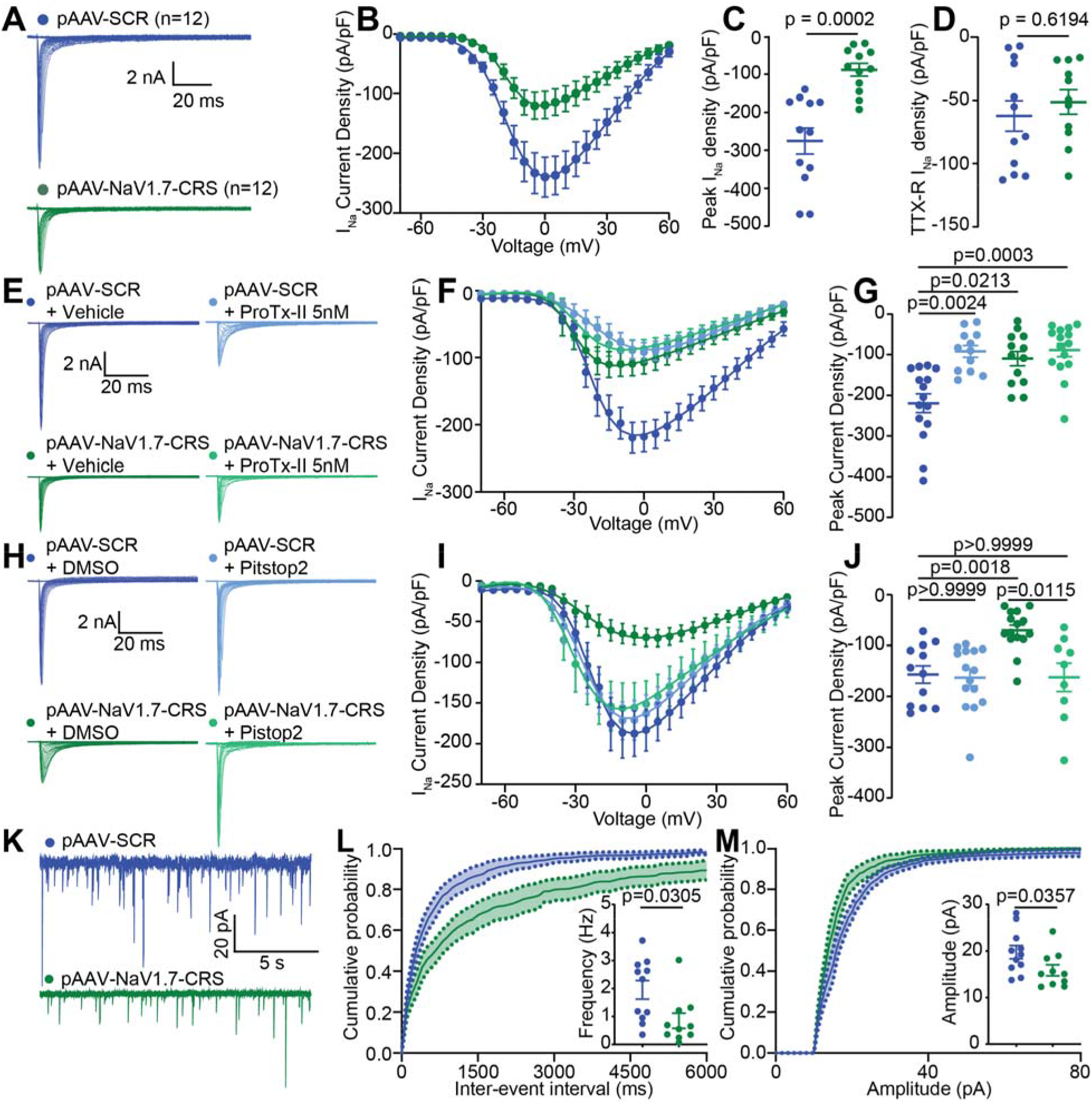
AAV plasmid encoding NaV1.7-CRS reduces NaV1.7 currents through an endocytic mechanism and decreases spontaneous activity in the spinal cord. (**A**) Representative current traces recorded from DRGs transfected with an AAV plasmid encoding a scrambled control peptide (pAAV-SCR, blue, n=12) or a peptide corresponding to the CRMP2 regulatory sequence of NaV1.7 (pAAV-NaV1.7-CRS, green, n=12). (**B**) Summary plots of the current density-voltage relationship for each condition fitted with a Boltzmann curve, (**C**) peak current density and (**D**) Electrically isolated TTX-R currents. (**E**) Representative current traces recorded from DRG neurons transfected as indicated andtreated with vehicle or 5nM of ProTx-II. (**F**) Summary plots of the current density-voltage relationship for each condition fitted with a Boltzmann curve and (**G**) Peak current density quantified from curves in panel E. (**H**) Representative current traces from groups transfected with either pAAV-SCR or pAAV-NaV1.7-CRS and treated with the clathrin-mediated endocytosis inhibitor Pitstop2 (20μM). (**I**) Summary plots of the current density-voltage relationship and (**J**) Peak current density for each experimental condition. Values for biophysical parameters are summarized in **Table S3**. n=9-15 cells (**K**) Representative traces of spontaneous excitatory postsynaptic currents (sEPSC) recordings from rat substantia gelatinosa (SG) neurons transduced with AAV-SCR (blue) or AAV-NaV1.7-CRS (green). (**L**) Cumulative distribution and bar graph of sEPSC inter-event intervals recorded from Substantia Gelatinosa (SG) neurons transduced with NaV1.7-CRS encoding plasmid (**M**) Cumulative distribution and bar graph showing decreased sEPSC amplitude in slices from AAV-NaV1.7-CRS treated rat compared to control. Data are expressed as means ± SEM. Mann-Whitney and kurskal wallis test (details in **Table S5**).

NaV1.7 regulates spinal nociceptive neurotransmission from the sensory neuron synapse to the second order neurons in the lamina I. We then tested if AAV-NaV1.7-CRS could silence spinal neurotransmission similar to NaV1.7 inhibition (*21, 35*). In spinal cord prepared from rats injected with AAV-NaV1.7-CRS (**Fig. 5K**), we recorded a decreased frequency (**Fig. 5L**) as well as decreased amplitude of spontaneous excitatory post-synaptic potentials (sEPSC) (**Fig. 5M**), when compared to AAV-SCR control. Collectively, these data show that NaV1.7 inhibition via targeting of the CRS domain recapitulates the spinal phenotype of NaV1.7 loss of function.

### AAV9 delivery of NaV1.7-CRS reverses and prevents the development of mechanical allodynia following SNI

Our in vitro observations demonstrated that competitive blockade of the CRS domain using either a peptide (**Fig. 2**) or a genetically encoded sequence (**Fig. 5**) leads to the complete inhibition of NaV1.7 functions in DRG sensory neurons. To test the long-term effects of blocking the CRS domain, we packaged the AAV-NaV1.7-CRS plasmid into a functional AAV9 capsid (**Fig. 6A**). Intrathecal injection (lumbar puncture) of 1×10^10^ viral particles successfully allowed for the expression of our construct in both DRG (**Fig. 6B**) and spinal cord (**Fig. 6C**). We next leveraged this gene therapy approach to test the potential tolerance and unwanted effects of constant NaV1.7 inhibition by CRS targeting in vivo (**Fig. S9**). Male mice with SNI were injected with AAV9-NaV1.7-CRS or control virus. Mice injected with AAV9-NaV1.7-CRS demonstrated a reversal of mechanical allodynia that was maintained for more than 30 days following the injection compared to control-injected mice (**Fig. 6D-E**). Before termination of the experiment, mice were subjected to an open field test wherein they exhibited no impairment of motor capacity (**Fig. 6F**) or developed any anxiety-like behaviors (**Fig. 6G**). The relief of mechanical allodynia and lack of other effects was recapitulated in female mice injected with AAV9-NaV1.7-CRS (**Fig. 6H-K**). These data show that targeting the CRS domain of NaV1.7 can provide complete and sustained relief of painful behaviors in male and female mice with chronic neuropathic pain.

**Figure 6.**
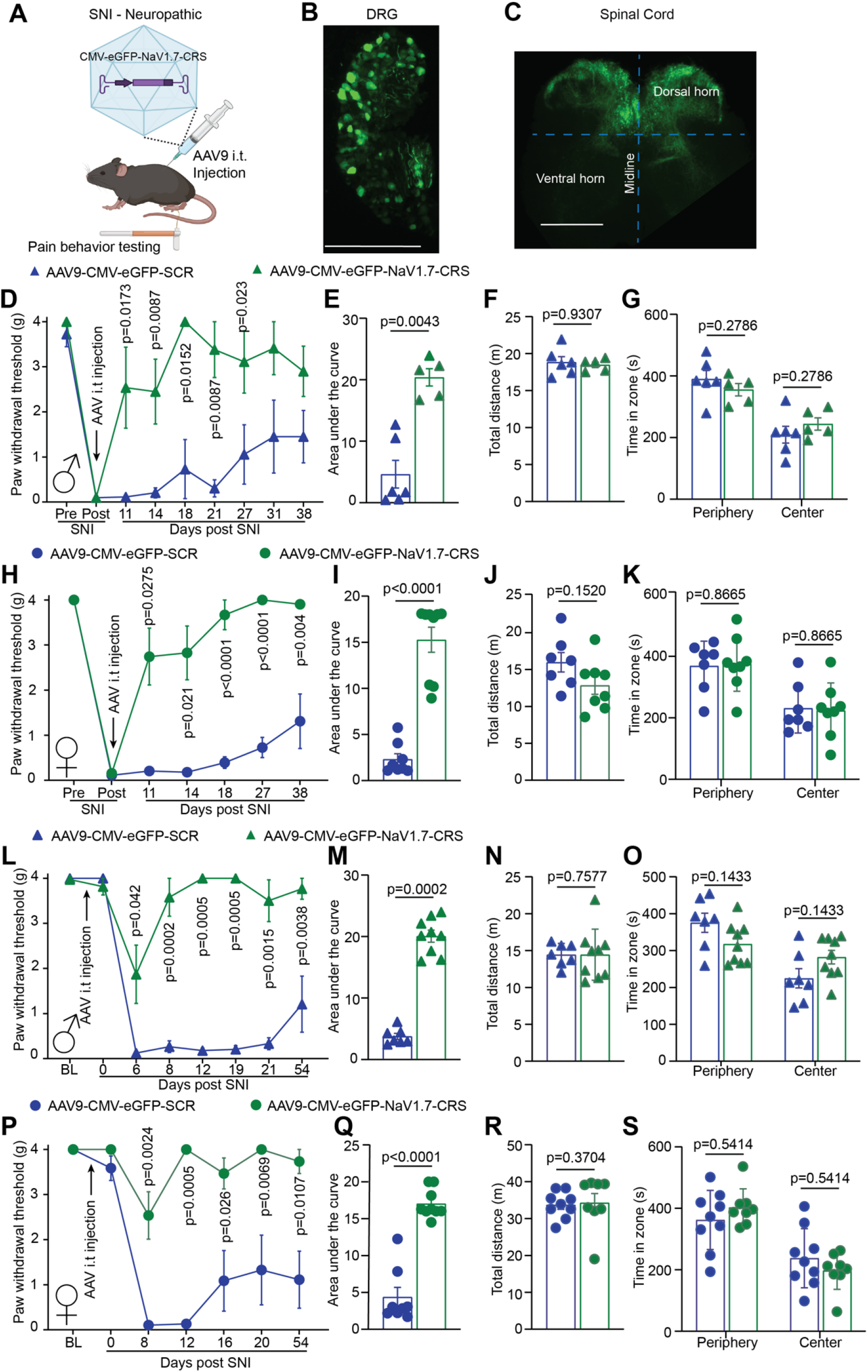
AAV9 plasmid encoding NaV1.7-CRS reverses and prevents mechanical allodynia in a mouse model of neuropathic pain. (**A**) Cartoon showing the experimental paradigm for testing an AAV9 vector to express the CRS domain in mice. Neuropathic pain was induced through the SNI model and virus was intrathecally delivered following development of neuropathic pain (see **Fig. S9**). Visualization of GFP fluorescence indicating successful injection of AAV9-NaV1.7-CRS in the DRGs (**B**) and in the spinal cord (**C**) of mice. (**D**) Paw withdrawal thresholds of male mice intrathecally injected with AAV9 virus encoding either NaV1.7-CRS or the scrambled control peptide 7 days after SNI. (**E**) Area under the curve showing efficient pain reversal by AAV-NaV1.7-CRS. (**F**) Evaluation of locomotor activity and (**G**) anxiety-like behavior using the open field test in mice treated as indicated. Data was replicated in female mice for paw withdrawal threshold (**H**), area under the curve (**I**), locomotor activity (**J**) and anxiety (**K**). Together showing that AAV9-NaV1.7-CRS reversed chronic pain in both male and female mice. Intrathecal injection of AAV9 was performed 7 days before SNI surgery (see **Fig. S9**). Paw withdrawal thresholds (**L**) and area under the curve (**M**) show the prevention of chronic allodynia in male mice injected with AAV9-NaV1.7-CRS. No effect was detected in locomotor activity (**N**) and anxiety-like (**O**) behaviors in the open field test. Data was replicated in female mice for paw withdrawal threshold (**P**), area under the curve (**Q**), locomotor activity (**R**) and anxiety (**S**). Together showing that AAV9-NaV1.7-CRS prevented chronic pain development in both male and female mice. n=5-9 animals; error bars indicate mean ± SEM; detailed statistical analysis is in **Table S5**. The experiments were conducted by investigators blinded to treatments.

As other transgenic models of NaV1.7 loss of function (*32, 33, 36*) were resistant to the development of chronic neuropathic pain, we next asked if targeting the CRS domain could be used to prevent the development of chronic pain following a nerve injury. Naïve male and female mice were injected with AAV9-NaV1.7-CRS before being subjected to a SNI surgery (**Fig. S9**). In male mice, no change of the basal mechanical threshold was detected due to the injection of AAV9-NaV1.7-CRS compared to AAV9-SCR control (**Fig. 6L**). However, mice injected with AAV9-NaV1.7-CRS were resistant to the development of mechanical allodynia after SNI surgery while AAV9-SCR control mice became hypersensitive to mechanical stimuli (**Fig. 6L-M**). As above, mice had no impairment of motor or anxiety behaviors (**Fig. 6N-O**). We observed no sex difference as female mice showed a similar result (**Fig. 6P-S**). Collectively, these findings reveal that the CRS domain can be targeted to prevent the initiation and maintenance of chronic neuropathic pain in both male and female mice. Therefore, the AAV9-NaV1.7-CRS could be used as a novel genetic therapy to provide for long-lasting pain relief.

### AAV9-NaV1.7-CRS reduces Na^+^ currents through NaV1.7 channels in macaque sensory neurons

To validate our findings in a translationally relevant model organism, we gained access to Rhesus macaque DRG and spinal cord tissues. We hybridized macaque spinal cord lysate on our peptide array and found that macaque CRMP2 bound to the same CRS sequence in the intracellular loop 1 of NaV1.7 (**Fig. S10A**). Next, by co-immunoprecipitation, we showed that (*i*) the CRMP2/NaV1.7 interaction was conserved from rodents to non-human primates and that (*ii*) the Myr-TAT-NaV1.7-CRS peptide could block this interaction in macaque spinal cord lysates (**Fig. S10B**). We then treated cultured macaque DRG neurons with the AAV9-NaV1.7-CRS and recorded sodium currents (**Fig. S10C**). Sensory neurons transduced with AAV9-NaV1.7-CRS had reduced total sodium current density (**Fig. S10D-E**) compared to the AAV9-SCR treated group. Additionally, treatment with the NaV1.7 direct blocker ProTx-II demonstrated that maximal inhibition was achieved following transduction with AAV9-NaV1.7-CRS (**Fig. S10D-E**). As before, AAV9-NaV1.7-CRS had no impact on the half maximal activation voltage (**Fig. S10F, Table S4)**). To test if TTX-R currents were impacted by AAV9-NaV1.7-CRS in macaque DRG neurons, we applied TTX. In this condition, AAV9-NaV1.7-CRS and AAV9-CRS control were indistinguishable (**Fig. S10G-J**). Altogether, our results prove that selective NaV1.7 inhibition can be achieved by targeting the CRS domain in DRG neurons from non-human primates.

## DISCUSSION

We report the discovery and characterization of a previously unknown intracellular domain of NaV1.7, necessary for the membrane localization and function of the channel. We validated this CRMP2 regulatory sequence in NaV1.7 across five different species from rodents (mice and rats) to more translational systems (pig, macaque and human). We further demonstrated that while the CRS domain is conserved across species, it is divergent among the NaV1.x subtypes. Inhibition of the CRS domain either with a decoy peptide or genetically (*i*) selectively decreased NaV1.7 current density in rat and macaque DRG neurons, (*ii*) dampened DRG neuron excitability via (*iii*) decreased membrane localization of NaV1.7, (*iv*) reduced spinal neurotransmission, which (*v*) attenuated neurotransmitter release from the spinal cord. Additionally, we demonstrated in vivo that competitive inhibition of the CRS domain with a blocking peptide reversed mechanical allodynia in male and female rats with chronic neuropathic pain. In mice, we used a gene therapy approach to show that CRS inhibition could prevent the initiation and maintenance of chronic neuropathic pain. These effects were achieved without any unwanted effects on physiological pain thresholds, motor coordination or anxiety. Our findings point to the identification of a targetable domain on NaV1.7 that can be leveraged to develop safe and selective pain-relieving drugs.

Our previous studies identified CRMP2 as a major regulator of NaV1.7 membrane localization. We reported that NaV1.7 internalization is regulated by formation of a complex between CRMP2, the endocytic adaptor Numb, the E3 ubiquitin ligase Nedd4-2, and the endocytic protein Eps15 (*19*). We developed a novel compound, **194**, which inhibits CRMP2 SUMOylation to selectively reduce NaV1.7 currents. **194** reversed mechanical allodynia in multiple animal models of neuropathic pain (*21, 37, 38*). However, the underlying basis of the selective regulation of NaV1.7 by CRMP2 was never elucidated. We now conclude that this specific regulation is achieved via a specific binding domain for CRMP2 on NaV1.7 that is not found in any other sodium channels. Using the TTX-R Halo-NaV1.7 construct allowed us to investigate the role of the CRS domain directly in DRG neurons by identifying transfected cells and eliminating the contribution of endogenous sodium channels. We thus found that the CRS domain on NaV1.7 is necessary for the membrane localization and function of the channel. We further show that competitive inhibition of the CRS domain either with a peptide or genetically can selectively inhibit NaV1.7. This is supported by the lack of further blockade of sodium currents by ProTx-II treatment of rat or macaque DRG neurons treated with Myr-TAT-NaV1.7-CRS but also by our observation that ProTx-II did not induce a rightward shift of half voltage activation in these neurons, thus showing that NaV1.7 was no longer available. Importantly, our data consistently supports clathrin-mediated internalization as the mechanism by which NaV1.7 can be downregulated to provide pain relief. Rather than blocking the pore of the channel directly, indirect modulation of NaV1.7 function by dissociating accessory proteins such as CRMP2, helped us to achieve a more specific inhibition of the channel. This was further validated in a screen against all orphan human GPCRs (PDSP) where the Myr-TAT-NaV1.7-CRS peptide had no effect.

The unique advantage of targeting the CRS domain may reside in its specific involvement in chronic neuropathic pain. CRMP2 extracted from the ipsilateral (painful) spinal dorsal horn of rats with SNI showed an enhanced binding to the CRS sequence. This is in line with our previous findings showing a specific involvement of CRMP2 in chronic but not in physiological pain thresholds. Accordingly, blocking the CRS sequence in vivo had no impact on acute pain models and on physiological pain thresholds in rats and in mice. However, we obtained an efficient reversal of mechanical allodynia in rodent models of chronic neuropathic pain. This indicates that strategies aiming at targeting the CRS domain of NaV1.7 will spare the physiological (protective) pain responses but silence pathological (chronic) pain.

Small molecule inhibitors have many advantages but have been notoriously difficult to develop for NaV1.7. A recent report described an alternative approach based on in-situ repression of NaV1.7 using AAV mediated delivery of zinc finger proteins or dCas9 which successfully produced antinociception in multiple mouse models of inflammatory and neuropathic pain (*39*). Gene therapy is particularly attractive for pain treatment because it would replace other pharmacological options that require daily dosing and equate to treat a chronic disease with a chronic treatment. Here, we leveraged our identification of this unique domain regulating NaV1.7 to use as a basis for a genetic therapy approach. Viral delivery of the NaV1.7-CRS peptide reversed established chronic neuropathic pain but was also able to prevent its development when given before an injury. Thus, suggesting that targeting the CRS sequence could be disease modifying and treat established chronic pain. We did not observe any signs of toxicity for the duration of our experiment (more than one month after viral injection). Our additional measures of motor behavior and anxiety suggests a strong safety profile for sustained inhibition of the CRS domain of NaV1.7. These observations are in line with those made with another gene therapy aiming at silencing NaV1.7 expression in the pain pathway (*39*).

The findings presented here are not without some limitations. One limitation is that we did not fully identify if accessory proteins other than CRMP2 could bind to the CRS domain of NaV1.7. Our CRS decoy peptide would chelate all these proteins similar to the inhibition of CRMP2 binding to NaV1.7 but these unknown partners may be additional important regulators of the channel. Related to this, we did not identify the protein complex mediating the internalization of NaV1.7 when the CRMP2/NaV1.7 interaction is blocked. We showed before that CRMP2 loss of SUMOylation recruits endocytic proteins to internalize NaV1.7 and that preventing CRMP2 SUMOylation could be used as a treatment for chronic pain (*17, 21, 36*). However, we are unsure of the exact proteins mediating the internalization of NaV1.7 when CRMP2 is unbound from the CRS domain. Another limitation is that our study primarily used rodent models of pain and some recent studies have shown that the pharmacology of NaV1.7 inhibitors can vary between rat and human sensory neurons. Despite these reported differences, our data show that the NaV1.7-CRS serves as the binding site for CRMP2 from rat, pig, macaque, and human lysates, which indicates that our observations in rodent tissues may be translational to humans. Finally, gene therapies have some potential drawbacks. Recent reports of neuronal toxicity were especially noted for the AAV9 serotype (*40*), which we used here. Thus, it is possible that some long-term sensory neuron toxicity could be observed beyond the duration of our experiment which was designed primarily to assess the potential tolerance to the sustained inhibition of the CRS domain of NaV1.7.

In summary, we discovered a key intracellular domain on NaV1.7, essential for the membrane localization and function of the channel. We demonstrate that this domain can be targeted to reduce the excitability of sensory neurons, spinal neurotransmission and abolish chronic neuropathic pain. We expect that the targeting of the CRS domain holds the key to developing future specific and safe therapeutics to treat chronic pain.

## Acknowledgments

We thank Drs. Frank Porreca and Amol Patwardhan for helpful discussions on this project.

## Author contributions

R.K. and A.M. developed the concept, designed experiments, and supervised all aspects of this project; A.M. performed the peptide array experiments; M.K. and S.P.M. performed computational and structural studies; A.M., K.G. and H.J.S. performed electrophysiology experiments, analyzed data, and wrote the manuscript; P.D. performed the biochemistry experiments and oversaw the calcium imaging studies; D.R., S.L. and A.C.R. performed electrophysiology experiments; L.F.M. performed the binding studies using MST; and C.L.M. and S.L. collected and analyzed the behavior data. All authors had the opportunity to discuss results and comment on the manuscript;

## Data and materials availability

All data is available in the main text, figures, and supplementary materials.

## Supplementary Materials

### Materials and Methods

#### Supplementary Figures

Fig. S1. CRMP2 binding to NaV1.X channel peptides analogous to NaV1.7-CRS

Fig. S2. Validation of expression for Halo-NaV1.7(1.X) plasmids

Fig. S3. Swapping the NaV1.7-CRMP2 regulatory segment with the analogous domain from NaV1.1-NaV1.9 channels reduces TTX-S currents but spares TTX-R currents.

Fig. S4. Treatment with Myr-TAT-NaV1.7-CRS reduces CRMP2 binding to peptide 140 in rat spinal cord lysate

Fig. S5. Myr-TAT-NaV1.7-CRS peptide does not affect voltage-gated potassium and calcium channels

Fig. S6. Myr-TAT-NaV1.7-CRS does not affect CRMP2 SUMOylation or phosphorylation.

Fig. S7. Sensory neuron AP waveform parameters are not affected by acute treatment with the Myr-TAT-NaV1.7-CRS peptide

Fig. S8. Disruption of coupling between CRMP2 and NaV1.7 does not produce effects in off-target acute pain assays

Fig. S9. Experimental timelines for assessment of mechanical allodynia, locomotor activity and anxiety-like behaviors from male and female mice with SNI.

Fig. S10. Macaque DRG neurons transduced by AAV9-NaV1.7-CRS show reduced NaV1.7 currents and no effect on TTX-R currents.

#### Supplementary Tables

Table S1. Biophysical properties of Halo-NaV1.7(WT) channels and Halo-NaV1.7(X.X) mutants.

Table S2. Sodium channel gating properties following treatment with Myr-TAT-NaV1.7-CRS or Myr-TAT-SCR control peptide.

Table S3. Biophysical properties of sodium currents following genetic delivery of interfering peptide in rat DRG neurons.

Table S4. Biophysical properties of sodium currents following genetic delivery of interfering peptide in macaque DRG neurons.

Table S5. Details of statistical comparisons.

