## Supplementary Materials for "Identification and targeting of a unique NaV1.7 domain driving chronic pain"

Title: A unique CRMP2 binding site on the NaV1.7 channel defines regulatory specificity and inhibits chronic neuropathic pain

**Authors:** Kimberly Gomez^1,2†^, Harrison J. Stratton^3†^, Paz Duran^1†^, Santiago Loya^1,2^, Cheng Tang^1,2^, Aida Calderon-Rivera^1,2^, Liberty François-Moutal^3^, May Khanna^1,2^, Cynthia L. Madura^3^, Shizhen Luo^3^, Dongzhi Ran^3^, Lisa Boinon^3^, Samantha Perez-Miller^1,2^, Aubin Moutal^4^, and Rajesh Khanna^1,2^*

**Affiliations:**

^1^Department of Molecular Pathobiology, College of Dentistry, New York University, New York, NY, 10010, USA.

^2^NYU Pain Research Center, 433 1^st^ Avenue Room 822 New York, NY 10010, USA.

^3^Department of Pharmacology, College of Medicine, The University of Arizona, Tucson, AZ, 85724 USA.

^4^Department of Pharmacology and Physiology, School of Medicine, St. Louis University, St. Louis, MO, 63104, USA.

^†^Contributed equally to this work

*To whom correspondence should be addressed:

Dr. Rajesh Khanna, Department of Molecular Pathobiology, College of Dentistry, New York University, 433 1^st^ Avenue Room 822 New York, NY 10010, USA. ORCID #: 0000-0002-9066-2969

Supplementary Materials and Methods:

**MATERIALS AND METHODS**

**Study Design**

This study was designed with the aim of elucidating the mechanism of CRMP2 dependent regulation of NaV1.7 in the context of chronic pain. We approached this question from a standpoint of translating our findings to human therapies and to this end we made the decision to include the human NaV1.7 channel as often as was feasible. To identify the interaction domain between CRMP2 and human NaV1.7 we printed the cytoplasmic loops of the channel on a peptide array. We probed this array with lysates from three species to ensure our findings were the same across species. We used biochemical approaches to validate the interaction in cultured cells and investigated the effects of disrupting this interaction on NaV1.7 trafficking and CRMP2 post-translational modifications. We then sought to identify the electrophysiological consequences of disrupting this interaction using DRG neurons transfected with a mutant NaV1.7 encoding plasmid or treated with an interfering peptide. Furthermore, we investigated the effects of our disruption strategy in multiple different pain models to assay the off-target and on-target actions of this approach. Sample sizes were determined based on our experience with electrophysiological, biochemical, and behavioral experiments in our laboratory. Experimenters were blind to the treatment and the animals were randomly assigned to experimental groups.

**Animals**

Adult male and female Sprague-Dawley rats (Pathogen-free male and female 100–250 g, Envigo, Placentia, CA) and male and female mice (C57BL/6NHsd, 20–24 g; Envigo) were kept in light (12-h light: 12-h dark cycle; lights on at 07:00 h) and temperature (23 ± 3°C) controlled rooms. Standard rodent chow and water were available *ad libitum.* All animal use was conducted in accordance with the National Institutes of Health guidelines, and the study was conducted in strict accordance with recommendations in the Guide for the Care and Use of Laboratory Animals of the University of Arizona (Protocol #: 16-141). Two adult male macaques (*Macaca mulatta*), aged 7 and 11 years, were used in this study. All animals were housed and bred in the University of Arizona Laboratory Animal Research Center. The same animals were tested using no more than 2 noninvasive behavioral assessments to reduce the number of mice and rats needed to complete the study. All efforts were made to minimize animal suffering. All behavioral experiments were performed by experimenters who were blinded to the genotype, and sex, and treatment groups.

**Materials and Reagents**

Peptides with the TAT cell penetrating sequence, YGRKKRRQRRR, fused to the CRMP2 binding domain of NaV1.7, SRGKCPPWWYRFAHK, were synthesized and HPLC-purified (>95% purity) by Genscript Inc. (Piscataway, NJ, USA). Scramble and random sequence-based peptides conjugated to various cargoes as controls have been previously studied as controls in molecular, biochemical, and behavioral assays and demonstrated to have no effects (*1-4*).

**Peptide SPOT Array**

Peptide spot arrays (15-mers with an overlap of 12 residues) spanning intracellular loops of human NaV1.7 were constructed using the SPOTS-synthesis method. Standard 9-fluorenylmethoxy carbonyl (Fmoc) chemistry was used to synthesize the peptides and spot them onto nitrocellulose membranes pre-derivatized with a polyethylene glycerol spacer (Intavis AG). Fmoc-protected and Fmoc-activated amino acids were spotted in 20 by 30 arrays on 150 mm by 100-mm membranes using an Intavis MultiPep robot. The nitrocellulose membrane containing the immobilized peptides was soaked in N-cyclohexyl-3-aminopropanesulfonic acid (CAPS) buffer (10 mM CAPS, pH 11.0 with 20% vol/vol methanol) for 30 minutes, washed once with Tris-buffered 0.1% Tween 20 (TBST), and then blocked for 1 hour at room temperature (RT) with gentle shaking in TBST containing 5% (mass/vol) nonfat milk and then incubated with rat, pig, and human spinal cord protein for 1 hour at RT with gentle shaking. Next, the membranes were incubated in primary antibody for CRMP2 for 2 hours at RT with gentle shaking, followed by washing with TBST. Finally, the membranes were incubated in secondary antibody (goat anti-rabbit DyLight 800, Cat# 355571, Thermo Fisher) for 45 minutes, washed for 30 minutes in TBST and visualized by infrared fluorescence (Li-Cor Odyssey Clx Imaging System; LI-COR Biosciences, Lincoln, NE). 2-4 independent peptide spot arrays were used in this study.

**Preparation of spinal cord lysates**

Lumbar segment of spinal cord lysates prepared from adult Sprague-Dawley rats were generated by homogenization and sonication in lysis buffer (50 mM Tris-HCl, pH 7.4, 50 mM NaCl, 2 mM MgCl_2_, 1% [vol/vol] NP40, 0.5% [mass/vol] sodium deoxycholate, 0.1% [mass/vol] sodium dodecyl sulfate [SDS]) as previously described (*5*). The lysis buffer included freshly added protease inhibitors (Cat# [B14002](https://www.ncbi.nlm.nih.gov/nuccore/B14002); Biotools, Houston, TX), phosphatase inhibitors (Cat# [B15002](https://www.ncbi.nlm.nih.gov/nuccore/B15002), Biotools), and Benzonase (Cat# 71206; Millipore, Billerica, MA). Protein concentrations were determined using the bicinchoninic acid protein assay (Cat# PI23225, Thermo scientific).

**Structural modeling and computational methods**

Secondary structure prediction was conducted using PSIPRED (*6*). Initially, the full 315 amino acid intracellular domain of NaV1.7 was submitted to SwissModel (*7*), Phyre2 intensive mode (*8*) and I-TASSER (*9*). SwissModel and Phyre2 returned homology models only for the peptide in the region covering amino acids 708-724 and 706-720, respectively, both consistent with a predominantly alpha helical structure. I-TASSER returned only low-quality models for the intracellular domain with the top results also consistent with a helical structure for the peptide. Subsequently, all 9 NaV isoform peptides were submitted to I-TASSER with their respective 6 flanking amino acids included at both ends (the equivalent of NaV1.7 amino acids 700-726), due to minimum size for prediction. The resultant models all exhibited a loop-helix for the equivalent of residues 706-720 with C-scores in the range of -1.82 to -0.64, estimated TM-scores from 0.49 to 0.63, and estimated RMSDs from 2.7 – 5.0 Å.

**Microscale Thermophoresis (MST)**

MST is an assay that measures the thermophoretic movement of molecules in optically generated microscopic temperature gradients allowing analysis of biomolecular interactions (*10*). In MST, increasing concentrations of unlabeled ligand are mixed with the fluorescently labeled biomolecule which is kept at a constant concentration. Purified CRMP2-His was fluorescently labelled with a His-Tag labeling kit RED-Tris-NTA (Nanotemper, Germany) per manufacturer’s instructions. Two hundred nanomolar of CRMP2-His (in PBS supplemented with 0.05% Tween-20 (PBS-T buffer)) was mixed with 1M NT-647-His-labeling dye. After incubating for 30 min at room temperature, labeled CRMP2 was pelleted by centrifugation at 15,000 x (g) for 10 min at 4°C. 50 nM of NTA-labeled His-CRMP2 was mixed with varying concentrations of the NaV1.7-CRS (SRQKCPPWWYRFAHK) or the analogous region from all other voltage gated sodium channels (NaV1.X) in PBS-T and incubated at room temperature for 10 min. The thermophoresis readouts were captured on a Monolith NT.115 (Nanotemper, Germany) using premium MST capillaries, at 40% LED and 40% MST power. Data analysis was performed with the MO Affinity Analysis software (Nanotemper) using the Kd model (standard fitting model derived from law of mass action).

**Catecholamine A Differentiated (CAD) Cell Culture**

Mouse neuron derived Cathecholamine A differentiated CAD cells (ECACC cat. no. 08100805, RRID: CVCL_0199) were grown in standard cell culture conditions, 37 °C in 5% (vol/vol) CO2. They were maintained in DMEM/F12 media supplemented with 10% (vol/vol) FBS (HyClone) and 1% penicillin/streptomycin sulfate from 10,000 μg/mL stock.

**Co-immunoprecipitation**

CAD cells were lysed into the IP buffer containing: 20 mM Tris·HCl, pH 7.4, 50 mM NaCl, 2 mM MgCl2, 10 mM N-Ethylmaleimide, 1% (vol/vol) Igepal CA-630, 0.5% (mass/vol) sodium deoxycholate, 0.1% (mass/vol) SDS with Protease inhibitors (cat. no. B14002; Biotool), and phosphatase inhibitors (cat. no. B15002, Biotool). Total protein concentration was determined by BCA protein assay (cat. no. PI23225; Thermo Fisher Scientific), and then 500 μg of total protein were incubated with 3 μg of CRMP2 antibody overnight at 4 °C under gentle agitation.

For Co-IP of endogenously SUMOylated proteins, 0.5% SDS was added to the lysates at 0.5% (mass/vol) final concentration, before boiling for 5 min at 95 °C. Then, 500 μg of total proteins were incubated with 5 μg of SUMO1 antibody overnight at 4 °C under gentle agitation. Protein G magnetic beads (cat. no. 10009D; Thermo Fisher Scientific), preequilibrated with the IP buffer, were then added to the lysates and incubated for 1 h at 4 °C to capture immunocomplexes. Beads were washed four times with IP buffer to remove nonspecific binding of proteins before resuspension in Laemmli buffer and boiling at 95 °C for 5 min before immunoblotting.

Identification of endogenous CRMP2 SUMOylation is challenging due to the presence of basal isopeptidase activity. To overcome this issue, we prepared CAD cell lysates in a denaturing buffer, followed by heat denaturing to inactivate SUMO isopeptidases and completely unfold the protein, which eliminates non-covalent SUMO1 interactions (*11*). We then performed immunoprecipitation with a SUMO1 antibody followed by western blot to detect SUMOylated CRMP2.

**CRMP2-HIS pull-down**
HIS tag Dynabeads (Cat# 10103D, Invitrogen), preincubated with purified CRMP2-HIS (0.5 µM), were incubated overnight with 300 µg of total protein from macaque spinal cord lysates at 4 ⁰C in the presence of the indicated peptides (5 µM) with gentle rotation. Beads were washed 3 times with lysis buffer before resuspension in Laemmli buffer and denaturation (5 minutes at 95 ⁰C) and immunoblotting as described previously.

**Cell-surface biotinylation**
Biotinylation was performed as described previously (*1, 12*). Briefly, live CAD cells were incubated with 0.5 mg·mL−1 sulfosuccinimidyl 6-(biotin-amido) hexanoate (EZ-Link Sulfo-NHS-LC-Biotin; cat. no. 21335; Thermo Fisher Scientific) for 30 min at 4 °C in cold PBS solution. Excess biotin was quenched by three washes with ice-cold PBS solution containing 100 mM glycine then washed three times with ice-cold PBS solution. The cells were lysed in lysis buffer containing: 20 mM Tris·HCl, pH 7.4, 50 mM NaCl, 2 mM MgCl2, 1% (vol/vol) Igepal CA-630 (cat. No. 19628; USBiological Life Sciences), 0.5% (mass/vol) sodium deoxycholate with protease inhibitors (cat. no. B14002; Biotool), and phosphatase inhibitors (cat. no. B15002; Biotool)]. The biotinylated proteins were separated by adsorption onto Dynabeads M-280 Streptavidin (cat. no. 11205D; Thermo Fisher Scientific) overnight at 4 °C. Beads were washed three times with lysis buffer, resuspended in Laemmli buffer, and heated at 95 °C for 5 min before immunoblotting.

**Immunoblotting**

Indicated samples were loaded on 4–20% Novex gels (cat. no. XP04205BOX; Thermo Fisher Scientific). Proteins were transferred for 1 h at 120 V using TGS [25 mM Tris, pH 8.5, 192 mM glycine, 0.1% (mass/vol) SDS], with 20% (vol/vol) methanol as transfer buffer to PVDF membranes (0.45 μm; cat. no. IPFL00010; Millipore), preactivated in pure methanol. After transfer, the membranes were blocked at room temperature for 1 h with TBST (50 mM Tris·HCl, pH 7.4, 150 mM NaCl, 0.1% Tween 20) with 5% (mass/vol) nonfat dry milk, and then incubated separately with the following primary antibodies diluted in TBST, 5% (mass/vol) BSA, overnight at 4 °C: NaV1.7 (cat. no. ab85015; Abcam), βIII-Tubulin (cat. no. G7121; Promega), CRMP2 (cat. no. C2993; Sigma-Aldrich), SUMO1 (cat. no. S8070; Sigma-Aldrich), CaV2.2 (cat. no. TA308673; Origene), Na+/K+ ATPase a-1 (cat. no. 05-369-25UG; Sigma-Aldrich), PSD-95 (cat. no. MA1-045; Invitrogen), Synaptophysin (cat. no. MAB5258; Sigma-Aldrich), CRMP2 pSer522 (cat. no. CP2191; ECM Biosciences) and CRMP2 pThr514 (cat. no. PA5-110113; Invitrogen). Next, the membranes were incubated in HRP-conjugated secondary antibodies from Jackson ImmunoResearch, and blots were developed by enhanced luminescence (WBKLS0500; Millipore) before exposure to photographic film.

**Calcitonin gene-related peptide (CGRP) release assay**
Rats were anesthetized with 5% isofluorane and then decapitated. Two vertebral incisions (cervical and lumbar) were made to expose the spinal cord. Pressure was applied to a saline-filled syringe inserted into the lumbar vertebral foramen, and the spinal cord was extracted. Only the lumbar region of the spinal cord was used for the CGRP release assay. Baseline treatments (#1 and #2) involved bathing the spinal cord in standard Tyrode solution. The excitatory solution, consisting of 90 mM KCl, was paired with the treatment for fraction #4. These fractions (5 minutes, 700 µL each) were collected for measurement of CGRP release. Samples were immediately stored in a −20˚C freezer. Myr-TAT-NaV1.7-CRS (5 µM), Myr-TAT-SCR, or vehicle (0.9% saline) was added to the pretreatment and cotreatment fractions (#3 and 4). The concentration of CGRP released into the buffer was measured by enzyme-linked immunosorbent assay (Cat# 589001; Cayman Chemical, Ann Arbor, MI).

**Synaptic fractionation and enrichment**

Adult rats were anesthetized using isoflurane and decapitated. Spinal cords were removed, and the dorsal horn of the spinal cord was dissected as this structure contains the synapses arising from the DRG. Synaptosomes isolation was done as described previously (*13*). Integrity of non-postsynaptic density (non-PSD) and PSD fractions was verified by immunoblotting. PSD95 was enriched in the PSD fractions while synaptophysin was enriched in non-PSD fractions. BCA protein assay was used to determine protein concentrations.

**Acute dissociation, culture, and transfection of dorsal root ganglia (DRG) neurons**
Dorsal root ganglia (DRG) were dissected from 100 g female Sprague-Dawley rats employing procedures described previously (*14*). For non-transfected cells, dissociated DRG neurons were plated onto 12 mm poly-D-lysine and laminin-coated glass coverslips and cultured for up to 24-48 h. For siRNA-Control and siRNA-CRMP2 transfection, collected cells were resuspended in Nucleofector transfection reagent containing siRNAs at a working concentration of 600 nM. Then, cells were subjected to electroporation protocol O-003 in an Amaxa Biosystem (Lonza), plated onto 12-mm poly-D-lysine- and laminin-coated glass coverslips and cultured for up to 48-72 h. siRNA transfection was verified by GFP fluorescence. Transfection efficiencies were between 20% and 30%, with approximately ∼10% cell death. For experiments with the Halo-NaV1.7 channels cells were transfected with 8µg of DNA before resuspension and plating onto glass coverslips. In experiments with the AAV plasmid encoding the NaV1.7 peptide or the scrambled control cells were mixed with 6 µg of DNA before transfection and plating.

**Calcium Imaging**

Changes in depolarization-induced calcium influx in rat DRG neurons were determined with Fura-2AM as previously described (*3, 15*). DRG neurons were incubated with 5 µM of Myr-TAT-peptides or DMSO as control, for a duration of 30 min. A standard bath solution containing 139 mM NaCl, 3 mM KCl, 0.8 mM MgCl2, 1.8 mM CaCl2, 10 mM Na-HEPES, pH 7.4, 5 mM glucose was used. Depolarization was evoked with a 10 sec pulse of 90 mM KCl.

**HaloTag and TTX-S current isolation**
This plasmid contains a HaloTag reporter enzyme that fluoresces in the presence of a cell penetrant Halo Ligand. To promote membrane localization of the channel and orient the HaloTag toward the extracellular membrane a portion of a β2 subunit was fused to the N-terminus. Importantly, this plasmid contains a mutation, Tyr-362 to Ser (Y362S), that confers resistance to the voltage gated sodium channel blocker tetrodotoxin (TTX) (*16, 17*). Furthermore, while the channel encoded by this plasmid is pharmacologically resistant to TTX, it retains the biophysical properties of NaV1.7. This is important because adult rat DRG neurons express tetrodotoxin-sensitive (TTX-S; mainly NaV1.1, NaV1.6 and NaV1.7) and tetrodotoxin-resistant (TTX-R; NaV1.8 and 1.9) voltage-gated sodium channels (*18, 19*). The kinetics of activation and inactivation of these channels is widely accepted to be different, with TTX-S channels exhibiting fast inactivation, and TTX-R channels displaying slower kinetics (*20*). We leverage the differences in biophysical properties of these channel populations to dissect TTX-R currents from TTX-S currents, which has been shown to be equivalent to pharmacological separation (*21*).

**Whole-cell voltage clamp electrophysiology**

Patch-clamp recordings were performed at room temperature (22–24°C). Currents were recorded using an EPC 10 Amplifier-HEKA linked to a computer with Patchmaster software. 5 µM of Myr-TAT-SCR and Myr-TAT-NaV1.7-CRS peptides and 10 µM of TAT-SCR and TAT-NaV1.7 peptides were applied acutely to the cells for ~ 10–30 minutes.

For total sodium current (I_Na_) recordings, the external solution contained (in mM): 130 NaCl, 3 KCl, 30 tetraethylammonium chloride, 1 CaCl2, 0.5 CdCl2, 1 MgCl2, 10 D-glucose and 10 HEPES (pH 7.3 adjusted with NaOH, and mOsm/L= 324). Patch pipettes were filled with an internal solution containing (in mM): 140 CsF, 1.1Cs-EGTA, 10 NaCl, and 15 HEPES (pH 7.3 adjusted with CsOH, and mOsm/L= 311). Peak Na^+^ current was acquired by applying 150-millisecond voltage steps from −70 to +60 mV in 5-mV increments from a holding potential of −60 mV to obtain the current-voltage (I-V) relation. Normalization of currents to each cell’s capacitance (pF) was performed to allow for collection of current density data. For I-V adjustments, functions were fitted to data using a non-linear least squares analysis (*22*). I-V curves were fitted using double Boltzmann functions:

*f = a+ g1/(1+exp((x-V_1/2_1)/k1)) + g2/(1+exp(-(x-V_1/2_2)/k2))*

where *x* is the prepulse potential, *V_1/2_* is the mid-point potential and *k* is the corresponding slope factor for single Boltzmann functions. Double Boltzmann fits were used to describe the shape of the curve, not to imply the existence of separate channel populations. Numbers *1* and *2* simply indicate first and second mid-points; *a* along with *g* are fitting parameters.

Activation curves were obtained from the I-V curves by dividing the peak current at each depolarizing step by the driving force according to the equation: *G = I/(V_mem_-**E_rev_)*, where *I* is the peak current, *V_mem_* is the membrane potential and *E_rev_* is the reversal potential. The conductance (G) was normalized against the maximum conductance (Gmax). Steady-state inactivation (SSI) curves were obtained by applying an H-infinity protocol that consisted of 1-second conditioning pre-pulses from −120 to +10 mV in 10-mV increments followed by a 200-millisecond test pulse to +10 mV. Inactivation curves were obtained by dividing the peak current recorded at the test pulse by the maximum current (Imax). Activation and SSI curves were fitted with the Boltzmann equation.

In experiments with the Halo-NaV1.7 channels cells were incubated with 200 nM of Halo Ligand Oregon Green for 2 minutes at 37^o^C. Following this labeling step cells were washed with fresh warm DMEM media 3x to remove unbound ligand. The cells were then left to recover for 30 minutes before recordings began. In all experiments where this channel was used recordings were performed in the presence of 500 nM TTX in the external solution to block endogenous NaV1.7 currents. During recording small-diameter DRG neurons successfully transfected with the Halo-NaV1.7 plasmid were identified using a fluorescence microscope with stimulation at 490 nm and emission at 520 nm.

In experiments in which clathrin assambly was inhibited, 20 µM of Pitstop2 was added to the cells 30 minutes prior the recordings. When NaV1.7-blocker was employed, Protox-II was added into the external recording solution at a final concentration of 5 nM.

To isolate potassium currents (I_K_), DRG neurons were bathed in external solution composed of (in millimolar): 140 N-methyl-glucamine chloride, 5 KCl, 1 MgCl2, 2 CaCl2, 10 D-glucose and 10 HEPES (pH adjusted to 7.3 with KOH and mOsm/L= 313). Recording pipettes were filled with internal solution containing (in mM): 140 KCl, 2.5 MgCl2, 4 Mg-ATP, 0.3 Na-GTP, 2.5 CaCl2, 5 EGTA, and 10 HEPES (pH adjusted to 7.3 with KOH and mOsm/L= 320). From a holding potential of -60 mV, IK activation was determined by applying 300-millisecond voltage steps from −80 to +60 mV in 10-mV increments.

Pipettes were pulled from standard wall borosilicate glass capillaries (Sutter Instruments) with a horizontal puller (Model P-97, Sutter Instruments). The input resistance of the pipettes ranged from 2 to 4 MΩ. Recordings were performed from small DRG neurons with capacitance between 10 and 35 pF (~18-33 μm). Series resistance under 7 MΩ was deemed acceptable. All experiments had a series resistance compensation between 60-90 %. Signals were filtered at 10 kHz and digitized at 10–20 kHz. Analyzes were performed by using Fitmaster software (HEKA) and Origin 9.0 software (OriginLab).

**Measurement of action potentials using whole-cell current-clamp electrophysiology**

For current-clamp recordings the external solution contained (in millimolar): 154 NaCl, 5.6 KCl, 2 CaCl2, 1 MgCl2, 10 D-Glucose, and 8 HEPES (pH 7.4 adjusted with KOH, and mOsm/L= 300). The internal solution was composed of (in millimolar): 137 KCl, 10 NaCl, 1 MgCl2, 1 EGTA, and 10 HEPES (pH 7.3 adjusted with KOH, and mOsm/L= 277). At room temperature (22–24°C), whole-cell patch clamp configuration was made, and current-clamp mode was performed to record action potentials. DRG neurons with a resting membrane potential (RMP) more hyperpolarized than −40 mV, stable baseline recordings, and evoked spikes that overshot 0 mV were used for experiments and analysis. The action potentials were evoked by current injection steps from 0–120 pA with an increment of 10 pA in 300 ms. Rheobase was measured by injecting currents from 0 pA with an increment of 10 pA in 50 ms. Analyses were performed by using Fitmaster software (HEKA) and Origin 9.0 software (OriginLab).

**Lumbar puncture for intrathecal treatment delivery**

Rats between postnatal day 12 and 15 (P12-15) were deeply anesthetized with 4% isoflurane for the induction of anesthesia and 2% for maintenance. The caudal half of the animal’s back was shaved and a 25G needle was inserted perpendicular to the spinal column through the L4-L5 intervertebral level and lowered until it met the vertebral body. Occasionally, a quick flicking of the tail could be observed and served as a proxy for optimal needle placement. Entry into the intrathecal space was indicated by a reduction in the pressure necessary to insert the needle. Animals were injected with 10 µL of indicated AAV9 virus containing either AAV-CMV-eGFP-NaV1.7-CRS or AAV-CMV-eGFP-SCR. Electrophysiological recordings were performed two days following the intrathecal injections to allow for cell mediated production of the plasmid encoded peptides.

**Preparation of spinal cord for whole-cell patch clam recordings**

Rats were deeply anesthetized with isoflurane (4% for induction and 2% for maintaining). For spinal nerve block, 0.3 mL of 2% lidocaine was injected to both sides of L4 to L5 lumbar vertebrae. Laminectomy was performed from mid-thoracic to low lumbar levels, and the spinal cord was quickly removed to cold modified ACSF oxygenated with 95% O2 and 5% CO2. The ACSF for rat dissection contained the following (in millimolar): 80 NaCl, 2.5 KCl, 1.25 NaH2PO4, 0.5 CaCl2.2H2O, 3.5 MgCl2.6H2O, 25 NaHCO3, 75 Sucrose, 1.3 ascorbate, 3.0 sodium pyruvate with pH at 7.4 and osmolarity at 310 mOsm. Transverse 400 µm-thick slices were obtained by a vibratome (VT1200S; Leica, Nussloch, Germany). Slices were then incubated for 45 mins at 37°C before a 1h incubation at RT in an oxygenated recording solution containing the following (in millimolar): 125 NaCl, 2.5 KCl, 1.25 NaH2PO4, 2 CaCl2.2H2O, 1 MgCl2.6H2O, 26 NaHCO3, 25 D-Glucose, 1.3 ascorbate, 3.0 sodium pyruvate with pH at 7.4 and osmolarity at 320 mOsm. The slices were then positioned in a recording chamber and continuously perfused with oxygenated recording solution at a rate of 3 to 4 mL/min before electrophysiological recordings at room temperature (RT).

**Whole-cell patch clamp electrophysiology of spinal cord slices**

Second order neurons located in lamina I/IIo of the Substantia Gelatinosa were visualized and identified in spinal cord slices using infrared differential interferences contrast video microscopy on an upright microscope (FN1; Nikon, Tokyo, Japan) equipped with a 3.40/0.80 water-immersion objective and near infrared (NIR) charged-coupled device camera (Pco Panda). Micropipettes with resistance of 6 to 10 MΩ when filled with internal solution were fabricated from borosilicate glass (1.5mm OD; Sutter Instruments, Novato, CA) with a four-step micropipette puller (P-97; Sutter Instruments, Novato, CA). The internal solution for recording spontaneous excitatory postsynaptic currents (sEPSC) contained the following (in mM): 120 potassium-gluconate, 20 KCl, 2 MgCl2.6H2O, 2.0 Na2-ATP, 0.5 Na-GTP, 20 HEPES, 0.5 EGTA with pH at 7.4 and osmolarity at 310 mOsm. In all recordings of spontaneous synaptic activity, whole-cell recording configuration was obtained in voltage-clamp mode and the membrane potential was clamped at -60 mV using a software controlled (PATCHMASTER, HEKA Elektronik, Lambrecht, Germany) digital patch clamp amplifier (EPC10; HEKA Elektronik, Lambrecht, Germany).

The whole-cell configuration was obtained in voltage-clamp mode. To record sEPSCs, bicuculline methiodide (10 µM, Cat# 14343, Sigma Aldrich) and strychnine (2 µM, Cat# S0532, Sigma Aldrich) were added to the recording solution to block γ-aminobutyric acid-activated (GABA) and glycine-activated currents. Hyperpolarizing step pulses (5 mV for 50 milliseconds) were periodically delivered to monitor the access resistance (15-25 MΩ), and recordings were discontinued if the access resistance deviated more than 20% from baseline values. sEPSCs were recorded from each neuron for a total duration of 2 minutes. Currents were sampled at 6 kHz and filtered at 3kHz then analyzed using the Mini-Analysis Program (Synatosoft Inc., NJ) to provide spreadsheets for the generation of cumulative probability plots. The frequency and amplitude of the recordings were compared between neurons from animals in the AAV-SCR and AAV-NaV1.7-CRS treated groups.

**AAV plasmid production and encapsulation of pAAV-NaV1.7-CRSin AAV9 capsid**
The plasmids pAAV-CMV-eGFP-NaV1.7 and pAAV-CMV-eGFP-SCR were generated by adding the NaV1.7-CRS peptide sequence and the scrambled peptide sequence into the AAV-CMV-eGFP core plasmid (plasmid #67634, Addgene). The sequences were cloned into the plasmid without the terminal lysine in frame following the GFP sequence. The following sequences were inserted into the plasmid: (NAV1.7) – AGCAGGCAGAAGTGCCCACCTTGGTGGTACAGATTCGCCCAC; (SCR) - AAGTACCACCCTTGGGCCTGCTTCAGGCAGTGGAGAAGCCCA. These plasmids were subcloned using the Clone EZ PCR cloning kit with high copy number and resistance to ampicillin (Genscript). Four micrograms of DNA were obtained for each plasmid from Genscript. The plasmids were subsequently packaged into AAV9 viral capsids to test the feasibility of disrupting the interaction between NaV1.7 and CRMP2 as a gene therapy. The encapsulation of the plasmid in AAV serotype 9 capsids was performed by the University of Arizona Viral Core facility. AAV9 particles were produced using HEK293T cells using the triple transfection method and purification of the particles was achieved using an iodixanol centrifugation gradient (*23*). Cells were transfected when approximately 80-90% confluent and the media was replaced with previously warmed media before transfection. The viruses were produced on 10 cm plates and each plate was transfected with 10 of pXR-capsid (pXR-9), 10 µg of recombinant transfer vector, and 10 µg of pHelper vector using polyethyleneimine at a ratio of 4:1. This mixture was incubated briefly at room temperature and then added in single drops to the cell culture media. After successful transfection, the virus was harvested 72-96h later and isolated using iodixanol density gradient ultracentrifugation (*24, 25*). The virus was then dialyzed with PBS supplemented with 50 mM NaCl and 0.00001% Pluronic F68 with 50-kDa filters to a final volume near 150 µL.

**Hot plate test**
Wild-type female and male mice were placed on a metal plate (Stoelting, Wood Dale, IL) warmed to 52°C and a timer was started. Latency to the first response (flinching or licking the hind paws or jumping) was recorded. A cutoff time of 10 seconds was used to prevent tissue damage, and any mouse that reached the cutoff time was designated the value of the cutoff time as their latency time.

**Tail-flick assay**

The distal third of the tail of WT female and male mice was immersed in warm water (52°C). Latency to remove the tail from the water (tail-flick latency) was recorded. To prevent tissue damage, a 10-second cutoff was used and animals that did not react before the cutoff were assigned a latency of 10 seconds

**Spared nerve injury (SNI) model of neuropathic pain and intrathecal catheter insertion**

The biceps femoris muscle of female and male rats was dissected to expose the 3 terminal branches of the sciatic nerve and perform spared nerve injury by transecting the common peroneal and tibial branches of the right sciatic nerve leaving the sural nerve intact (*26*). Before any testing, animals were allowed to recover for 5–7 days. Next, rats were sedated using a xylazine/ketamine/ (12/80 mg/kg intraperitoneal; Sigma-Aldrich) and placed in a stereotaxic head holder. Following the exposition and incision of the cisterna magna, an 8-cm catheter (PE-10; Stoelting) was implanted as previously described (*27*). The von Frey test was performed on the lateral side of the right plantar surface of the rats’ hind paws as described by Chaplan et al. (*28*), before and after 20µg/5µl of Myr-TAT-SCR and Myr-TAT-NaV1.7-CRS peptides were intrathecally administered.

**Assessment of motor coordination with the Rotarod assay**
Following placement of the intrathecal catheters, the rats were trained to walk on a rotating rod (8 rev/min; Rotamex 4/8 device) with a maximal cutoff time of 180 seconds. Training was initiated by placing the rats on a rotating rod and allowing them to walk on the rotating rod until they either fell off or 180 seconds was reached. This process was repeated 6 times and the rats were allowed to recover for 24 hours before beginning the treatment session. Prior to treatment, the rats were run once on a moving rod to establish a baseline value. Treatment, either Myr-TAT-NaV1.7-CRS or Myr-TAT-SCR, was administered spinally via the intrathecal catheter. Assessment consisted of placing the rats on the moving rod and timing until either they fell off or reached a maximum of 180 seconds. This was repeated 30 min after injection and then every hour for a total time course of 5 hours.

**Algogram**
The analgesic profiles of Myr-TAT-NaV1.7-CRS were analyzed by an in vivo screening tool, ALGOGramTM (ANS Biotech, Riom, France). This platform allowed us to obtain information about the effects of these peptides in 5 different pain areas (Acute and tonic pain, inflammatory pain, neuropathic pain, postoperative pain and visceral pain), by comparing their activity on a battery of 10 validated behavioral pain models with an ANS Biotech reference historical database. Assessment of the efficacy, and analgesic effects of a single administration (4 mg/ml, i.t.) of the peptides to adult male rats were analyzed in the rat models of: Tail flick test in healthy rats, paw pressure test in healthy rats, acetic acid-induced writhing, formalin test, Bennett model of peripheral mononeuropathy, oxaliplatin-induced neuropathy, carrageenan-induced mechanical hyperalgesia, kaolin-induced arthritis, Brennan model of incisional pain; and trinitrobenzene sulfonic acid (TNBS)-induced visceral hypersensitivity.

**Statistical methods and data analysis**

Graphing and statistical analysis was undertaken with GraphPad Prism (Version 9). All data sets were checked for normality using D’Agostino & Pearson test. Details of statistical tests, significance and sample sizes are reported in the appropriate figure legends. All data plotted represent mean ± SEM. For western blot experiments, statistical differences between groups were determined by Kruskal-Wallis test followed by the Dunn post hoc test. For electrophysiological recordings: peak current density was compared using Mann-Whitney tests, One-way ANOVA with the Tukey post hoc test and Kruskal–Wallis test with Dunnett’s post hoc comparisons; *V_1/2_* midpoint potential and *k* slope factor, were analyzed with Mann-Whitney test and one-way ANOVA with Tukey post hoc test; for sensory neuron excitability and action potential waveform parameters, statistical differences between groups were determined using multiple Mann-Whitney test and Mann-Whitney test. Statistical significance of thermal sensitivity was compared by Kruskal-Wallis test followed by the Dunn post hoc test. Behavioral data with a time course were analyzed by Multiple Mann-Whitney test and AUC was analyzed by Mann-Whitney test. For iCGRP release experiments, two-way ANOVA with the Sidak post hoc test was used to determine significance. Motor function differences were analyzed with Multiple Mann-Whitney test.

**Supplementary figures and legends.**

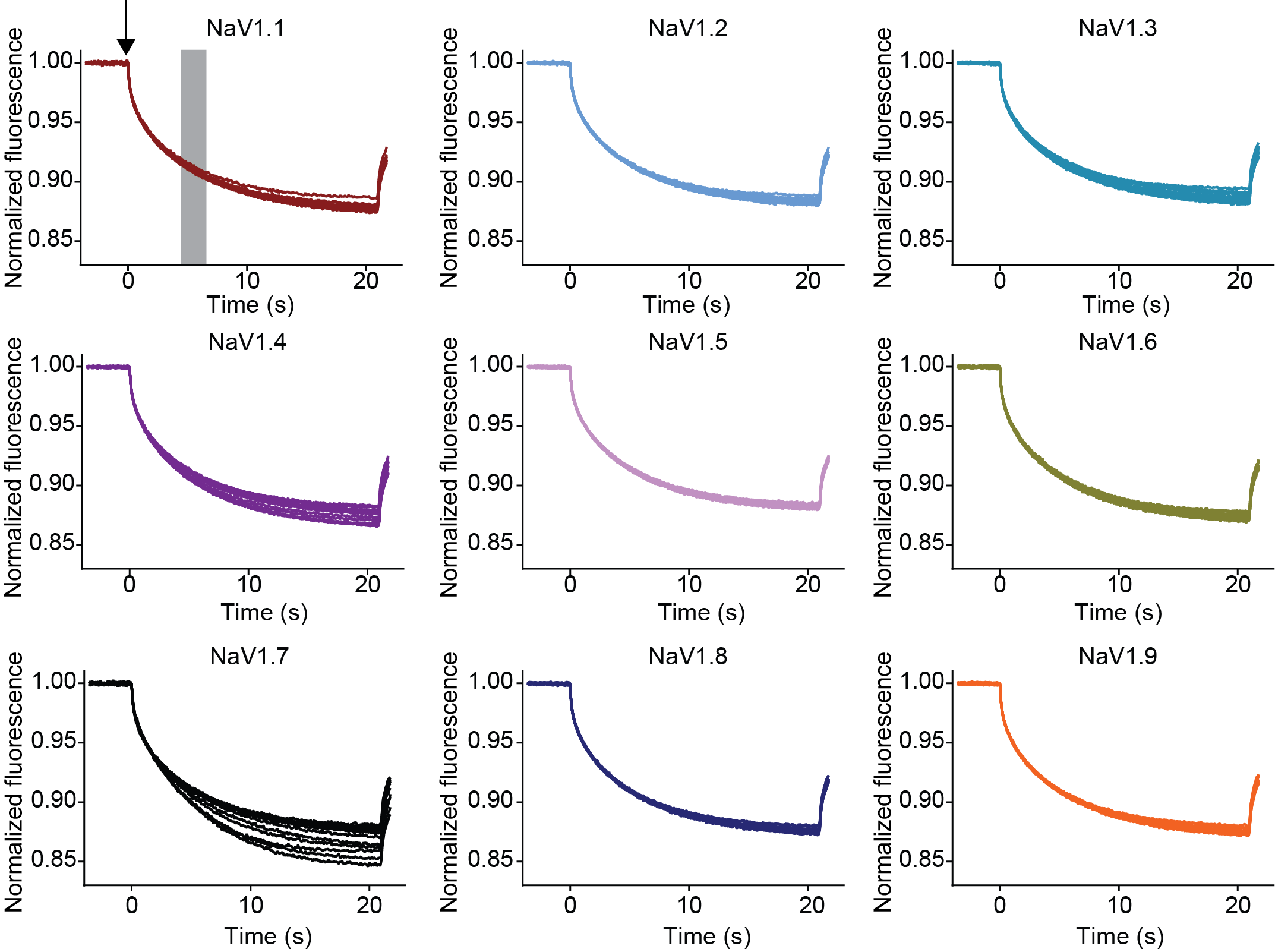

**Supplemental figure 1.** **CRMP2 binding to NaV1.X channel peptides analogous to NaV1.7-CRS.** Raw sensorgrams obtained from microscale thermophoresis experiments to determine binding between CRMP2 and peptides derived from intracellular loop 1 of NaV1.x channels, corresponding to data shown in Figure **1C**. The heating element of the thermophoresis began at the point in time indicated by the arrow. The gray box indicates the region of the curve used for fitting binding parameters.

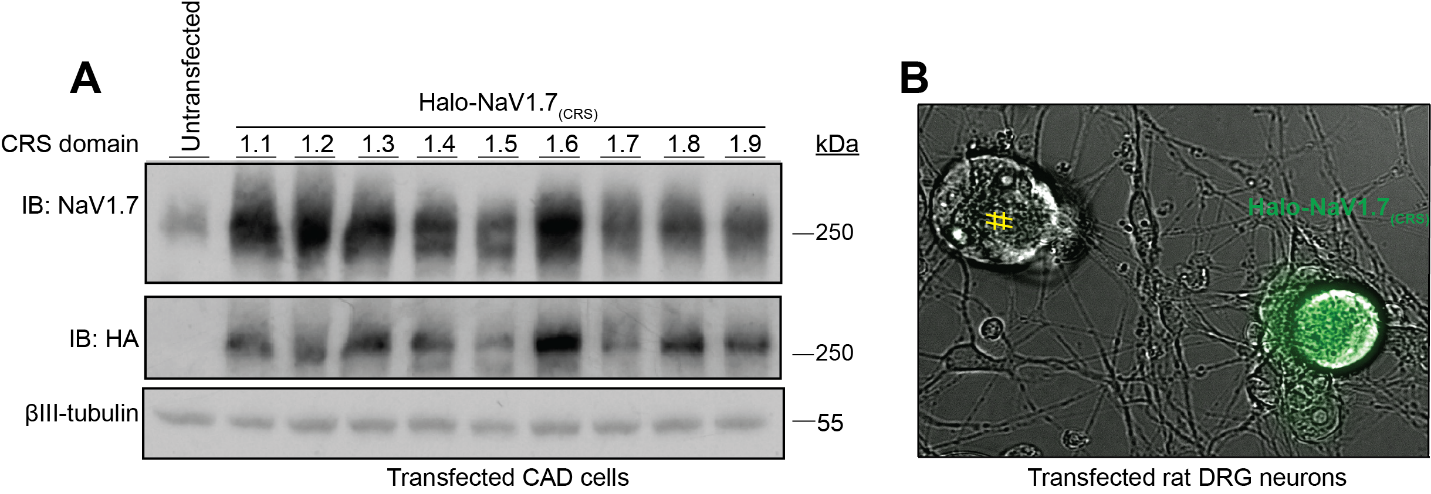
**Supplementary figure 2**. **Validation of expression for Halo-NaV1.7(1.x) plasmids**. (**A**) Western blots were performed after transfecting each Halo-NaV1.7(1.x) plasmid into CAD cells to verify the expression of channels in a cellular context. Each channel was constructed with an HA tag immediately adjacent to the HaloTag sequence, which allowed for identification of the overexpressed constructs separately from the endogenously expressed channels found in CAD cells. (**B**) DRGs from female rats transfected with the Halo-NaV1.7_(WT)_ channel and labeled with Oregon Green Halo Ligand. Note the non-transfected neuron labeled with the yellow pound sign.

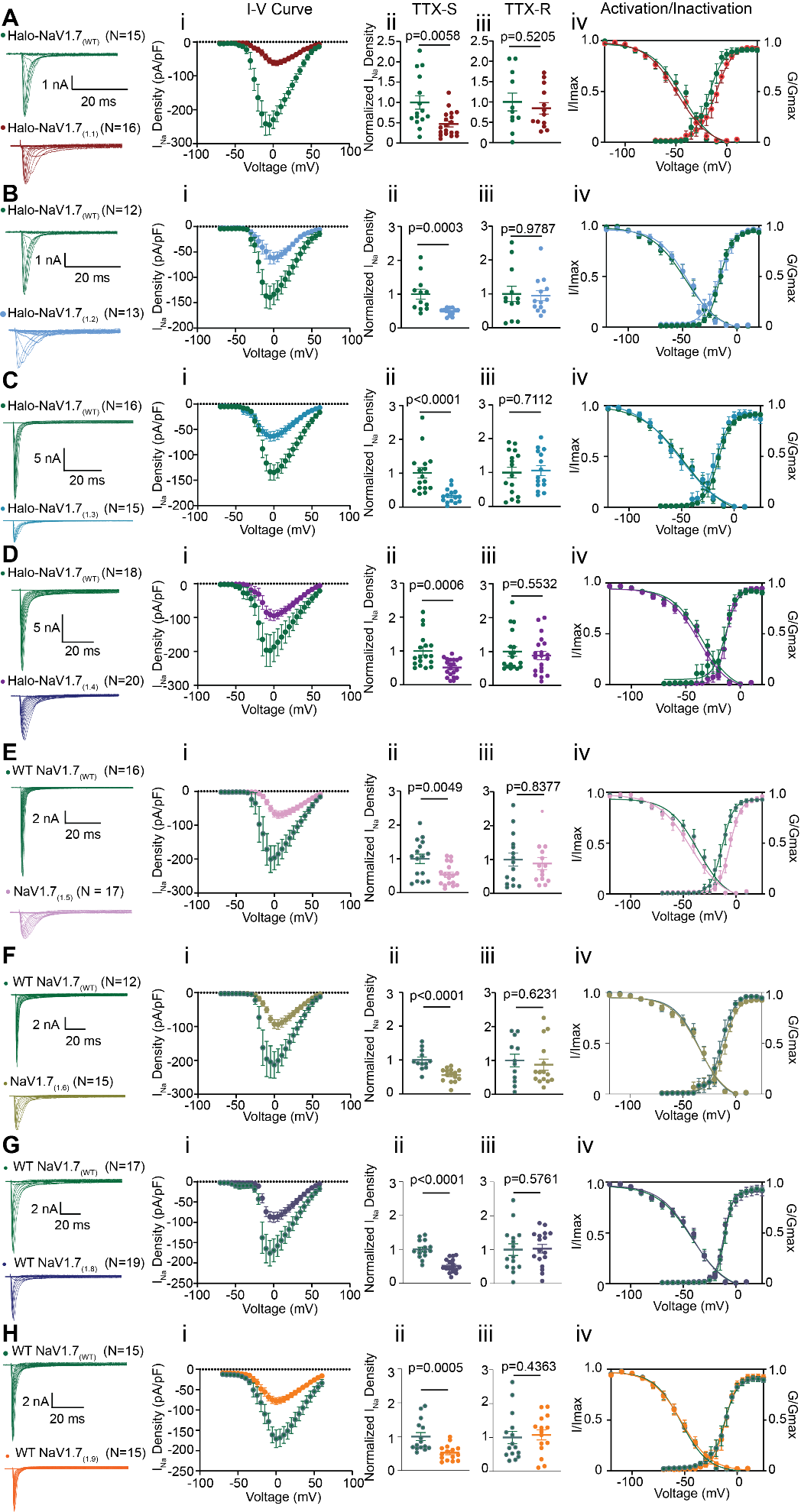

**Supplementary figure 3.** **Swapping the NaV1.7-CRMP2 regulatory segment with the analogous domain from NaV1.1-NaV1.9 channels reduces TTX-S currents but spares TTX-R currents.** Representative traces of sodium currents recorded from DRG sensory neurons transfected with Halo-NaV1.7_(WT)_ and (**A**) Halo-NaV1.7_(1.1_), (**B**) Halo-NaV1.7_(1.2)_, (**C**) Halo-NaV1.7_(1.3)_, (**D**) Halo-NaV1.7_(1.4)_, (**E**) Halo-NaV1.7_(1.5_), (**F**) Halo-NaV1.7_(1.6)_, (**G**) Halo-NaV1.7_(1.8)_, and (**H**) Halo-NaV1.7_(1.9_). (*i*) Current density-voltage relationship for the indicated Halo-NaV1.7 channels. (*ii*) TTX-S peak current density for the indicated Halo-NaV1.7_(1.x)_ channel compared to its own Halo-NaV1.7_(WT)_ control. (*iii*) TTX-R peak current density for the indicated Halo-NaV1.7_(1.x)_ channel compared Halo-NaV1.7_(WT)_, (*iv*) Biophysical properties of voltage-dependent activation and steady-state inactivation of the previously indicated channels. Values for *V_1/2_* and slope (*k*) can be found in Table 1; N=12-20 cells; error bars indicate mean ± SEM; *p* values as indicated; Mann-Whitney test.

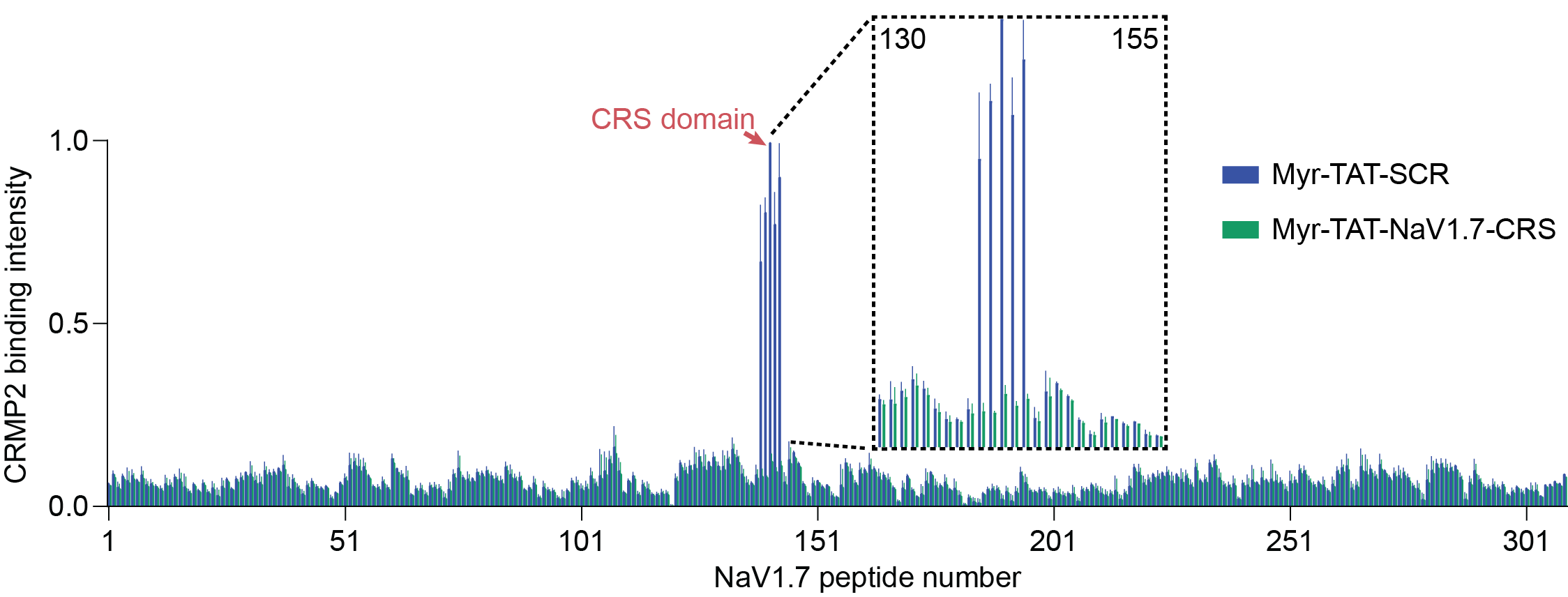

**Supplementary figure 4.** **Treatment with Myr-TAT-NaV1.7-CRS reduces CRMP2 binding to the CRS domain in rat spinal cord lysate.** Spinal cord lysates from rats were incubated with Myr-TAT-NaV1.7-CRS or Myr-TAT-SCR and binding to the peptide microarray was assessed. Treatment with Myr-TAT-NaV1.7-CRS (5 µM) significantly reduced the binding between CRMP2 and the CRS domain on the microarray.

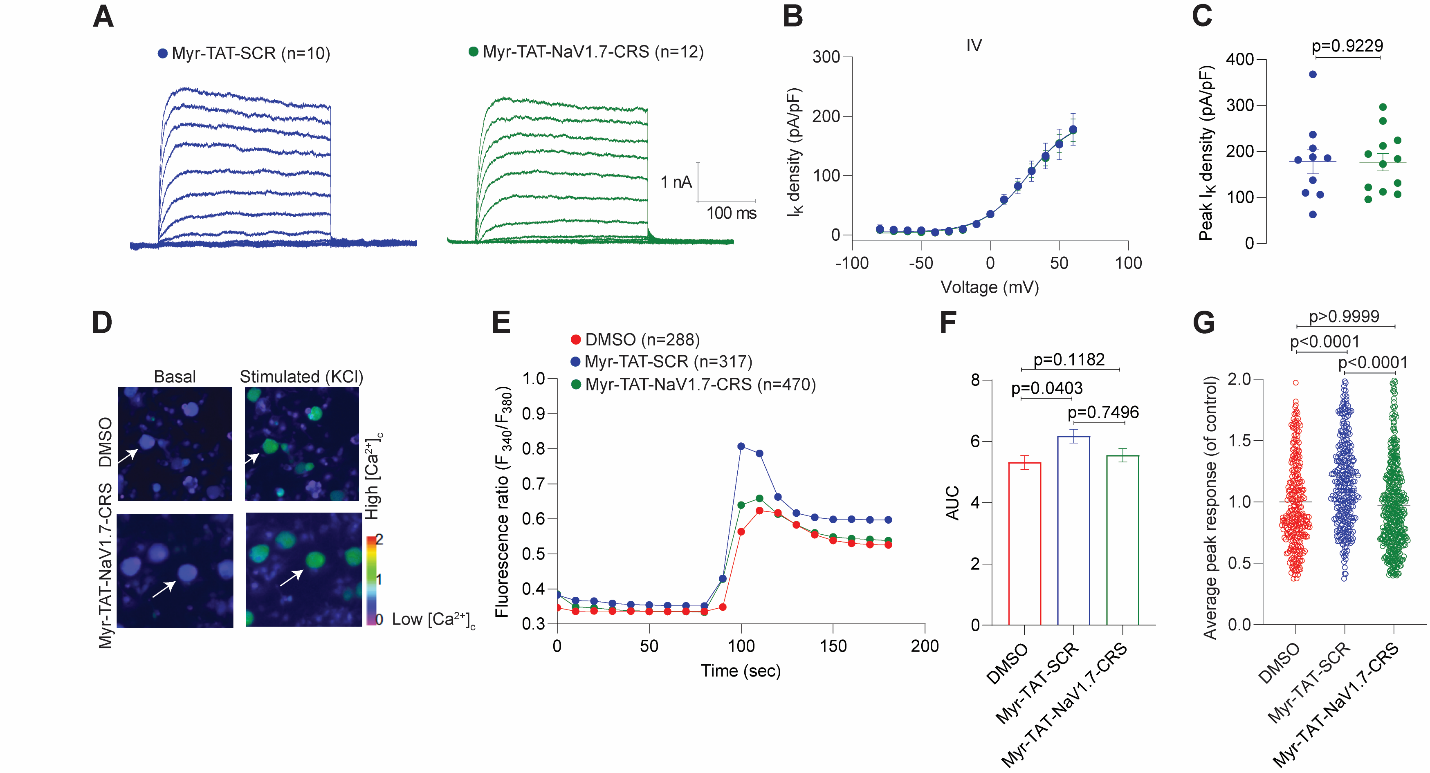

**Supplementary figure 5.** **Myr-TAT-NaV1.7-CRS peptide does not affect voltage-gated potassium and calcium channels.** (**A**) Representative potassium current traces recorded from sensory neurons in the presence of 5 µM of Myr-TAT-SCR and Myr-TAT-NaV1.7-CRS peptides. (**B**) Double Boltzmann fits for current density-voltage curves. (C) Summary of peak current densities (pA/pF). N=10-12 cells; error bars indicate mean ± SEM; *p* values as indicated; Mann-Whitney test. (**D**) Pseudo colored fluorescent images of a field of DRG neurons visualized for Fura-2AM, before (Basal) and after stimulation with 90 mM KCl for control (0.05% of DMSO) and 5 µM Myr-TAT-NaV1.7-CRS peptides treated neurons. Following a 1-min baseline measurement, neurons were stimulated with 90 mM KCl for 10 s. Exemplar images of the neurons (white arrows) at basal and peak calcium are indicated. (**E**) The average change in fluorescence ratio (F_340_/F_380_) over time for vehicle-treated (red circles), Myr-TAT-SCR treated (blue circles) or Myr-TAT-NaV1.7-CRS treated (green circles) neurons; all error bars are smaller than the symbols. Summary graph of the average area under the curve (AUC) between 80 and 180 s (**F**) and the peak fluorescence response (adjusted for background) (**G**) of rat neurons treated as indicated. N=288-470 cells; error bars indicate mean ± SEM; *p* values as indicated; One-way ANOVA with the Tukey post hoc test (**F**) and Kruskal-Wallis test with the Dunn post hoc test (**G**).

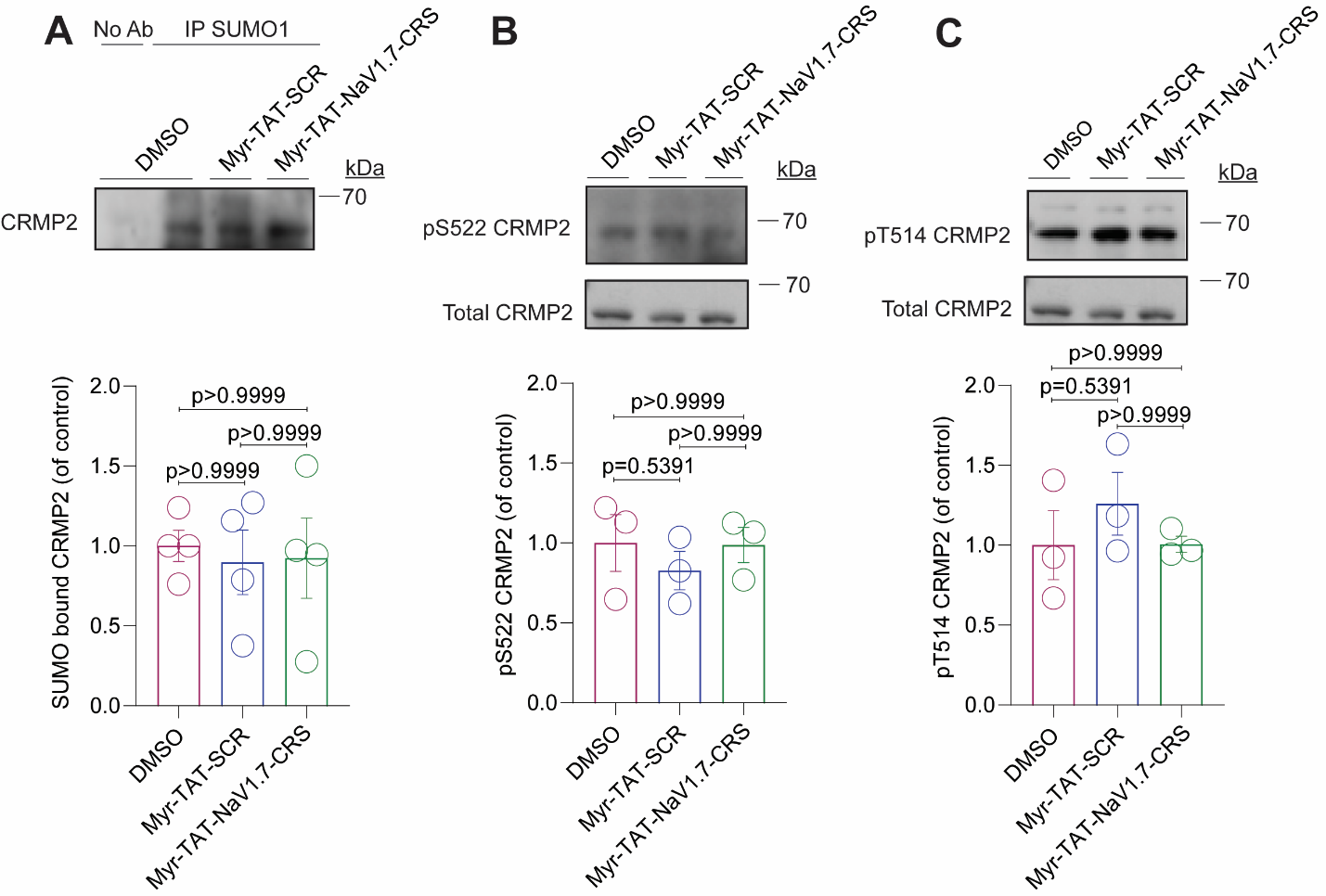

**Supplementary figure 6.** **Myr-TAT-NaV1.7-CRS does not affect CRMP2 SUMOylation or phosphorylation.** (**A**) Representative immunoblot (Top) and summary (Bottom) to detect SUMOylated CRMP2 from CAD cells treated with indicated peptides (n=4). (**B, C**) Representative immunoblots of CAD cells lysates indicating CRMP2 phosphorylation levels at indicated kinase target sites (n=3; Top), and quantitative analysis of CRMP2 phosphorylation compared with total CRMP2 (Bottom). Error bars show mean ± SEM; *p* values as indicated; Kruskal-Wallis test followed by the Dunn post hoc test.

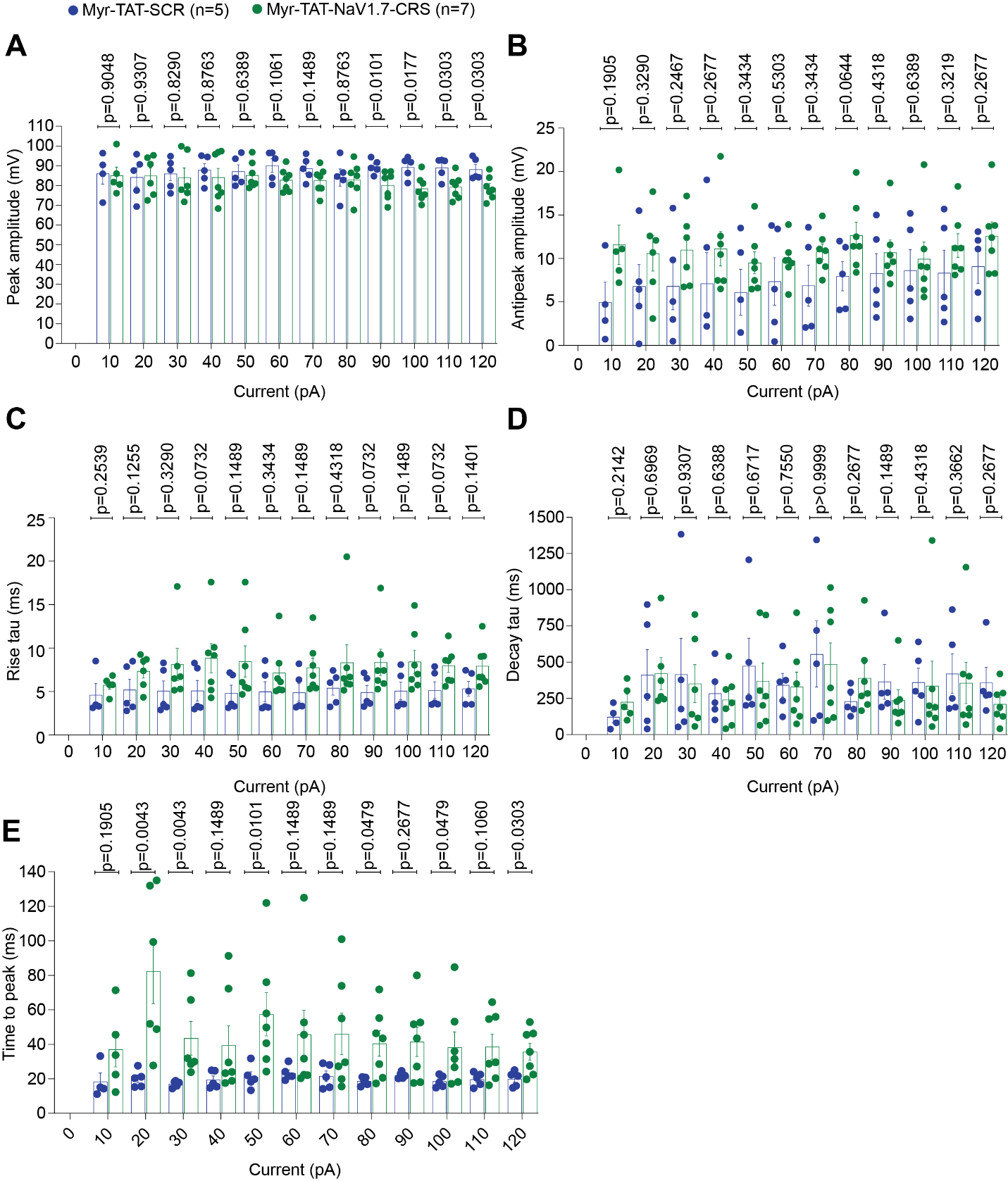

**Supplementary figure 7.** **Sensory neuron** **AP waveform parameters are not affected by acute treatment with the Myr-TAT-NaV1.7-CRS peptide**. Summary of the peak amplitude (in mV) (**A**), antipeak amplitude (in mV) (**B**), rise tau (in ms) (**C**), decay tau (in ms) (**D**) and time to peak (**E**) after incubation with 5 µM of Myr-TAT-SCR or Myr-TAT-NaV1.7-CRS peptides. N=5-7 cells; error bars indicate mean ± SEM; *p* values as indicated; Multiple Mann-Whitney test.

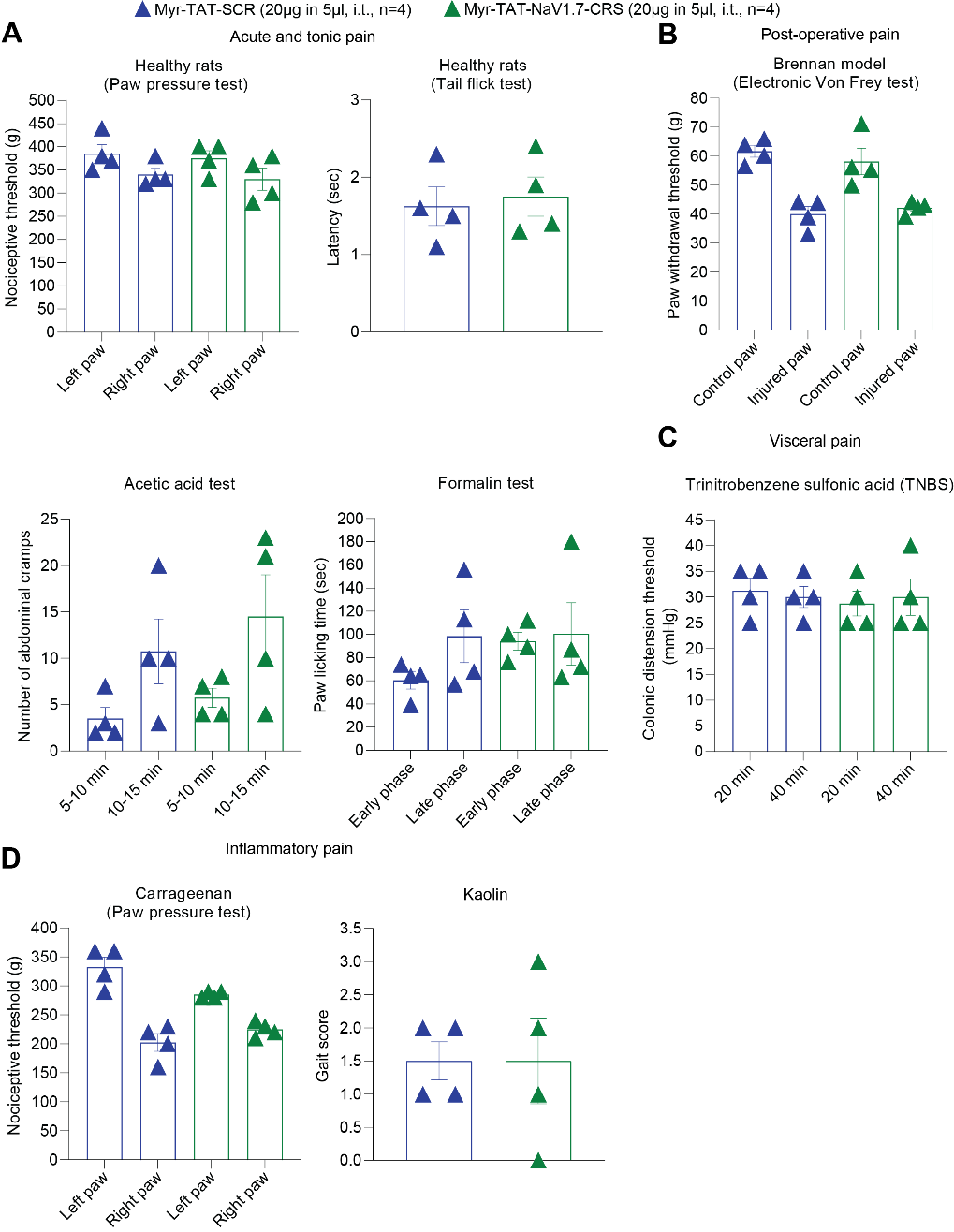

**Supplementary figure 8.** **Disruption of coupling between CRMP2 and NaV1.7 does not produce effects in off-target acute pain assays.** All behavioral assays were performed in male rats following a single 20µg in 5µl i.t. dose of either Myr-TAT-SCR (blue, n=4) or Myr-TAT-NaV1.7-CRS (green, n=4). (**A**) Top: Paw pressure test and the tail flick test demonstrated no effect on acute and tonic pain in healthy rats. Bottom: The number of abdominal cramps recorded for each group was not different between treatment categories for the acetic acid test. In the formalin test there were no differences observed between treatment groups in either the early or late phase as measured by paw licking time. (**B**) The Brennan model of post-surgical pain and (**C**) the model of trinitrobenzene sulfonic acid (TNBS) induced visceral pain. (**D**) Carrageenan and Kaolin were applied to the plantar surface of the paw and nociceptive thresholds and a gate score were determined. No effect was observed between groups treated with either peptide. Error bars indicated ± SEM; p values as indicated; Significant differences were determined with a Mann-Whitney.

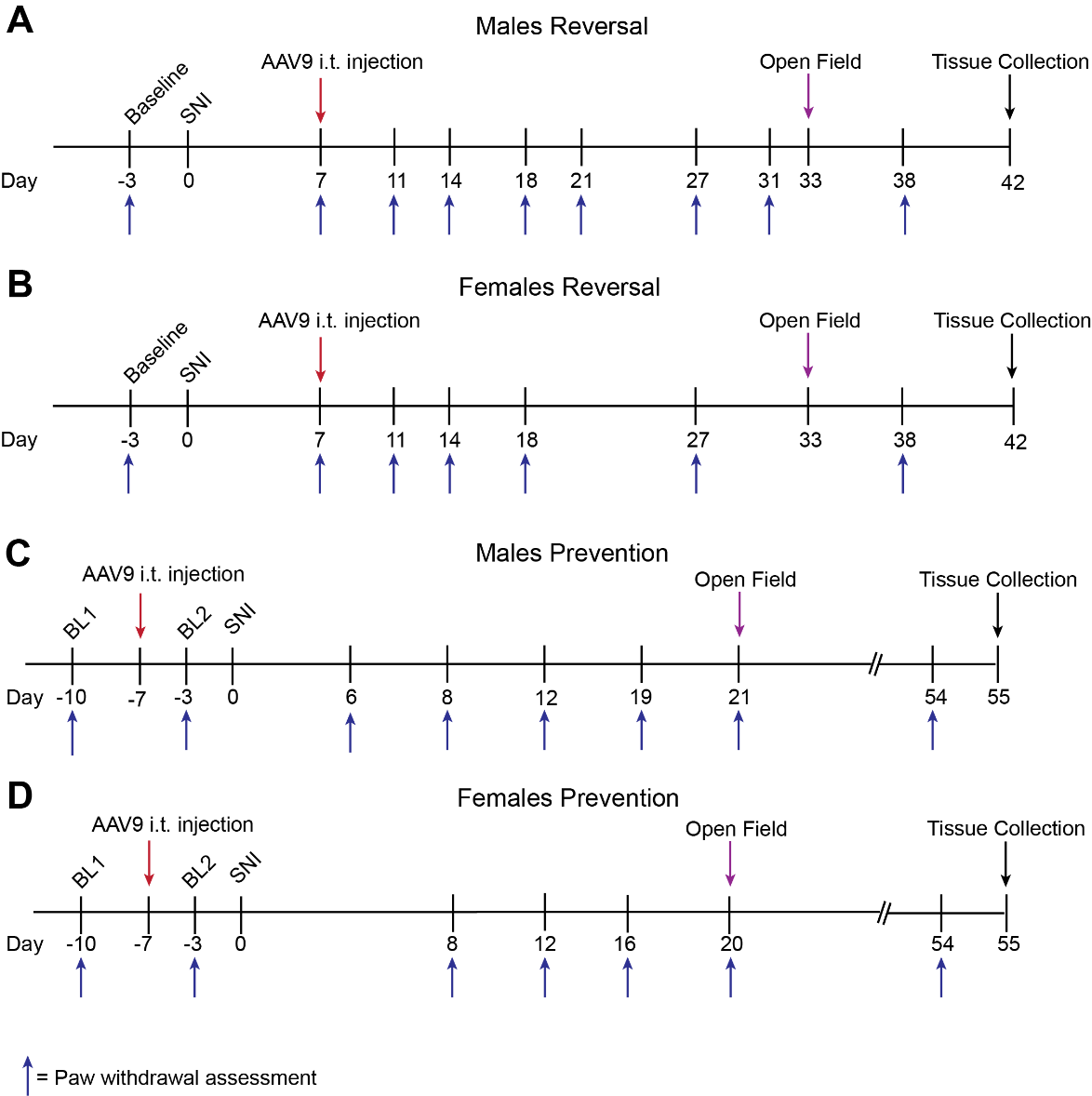

**Supplementary figure 9. Experimental timelines of paw withdrawal assessment, locomotor activity and anxiety-like behaviors from male and female mice with SNI.** (**A**) Male reversal timeline indicating assessment of mechanical allodynia at days -3, 7, 11, 14, 18, 21, 27, 31, and 38. AAV9 was intrathecally injected 7 days after SNI. Open field test was performed at day 33 and tissue was collected at day 42 after SNI. (**B**) Female reversal timeline indicating assessment of mechanical allodynia at days -3, 7, 11, 14, 18, 27, and 38. AAV9 was intrathecally injected 7 days after SNI. Open field test was performed at day 33 and tissue was collected 42 days after SNI. (**C**) Male prevention timeline indicating assessment of paw withdrawal at days -10, -3, 6, 8, 12, 19, 21, and 54. AAV9 was injected 7 days before SNI. Open field test was performed at day 21 and tissue was collected at day 55 after SNI. (**D**) Female prevention timeline indicating assessment of paw withdrawal at days -10, -3, 8, 12, 16, 20, and 54. AAV9 was injected 7 days before SNI. Open field test was performed at day 20 after SNI and tissue collection at day 55 after SNI.

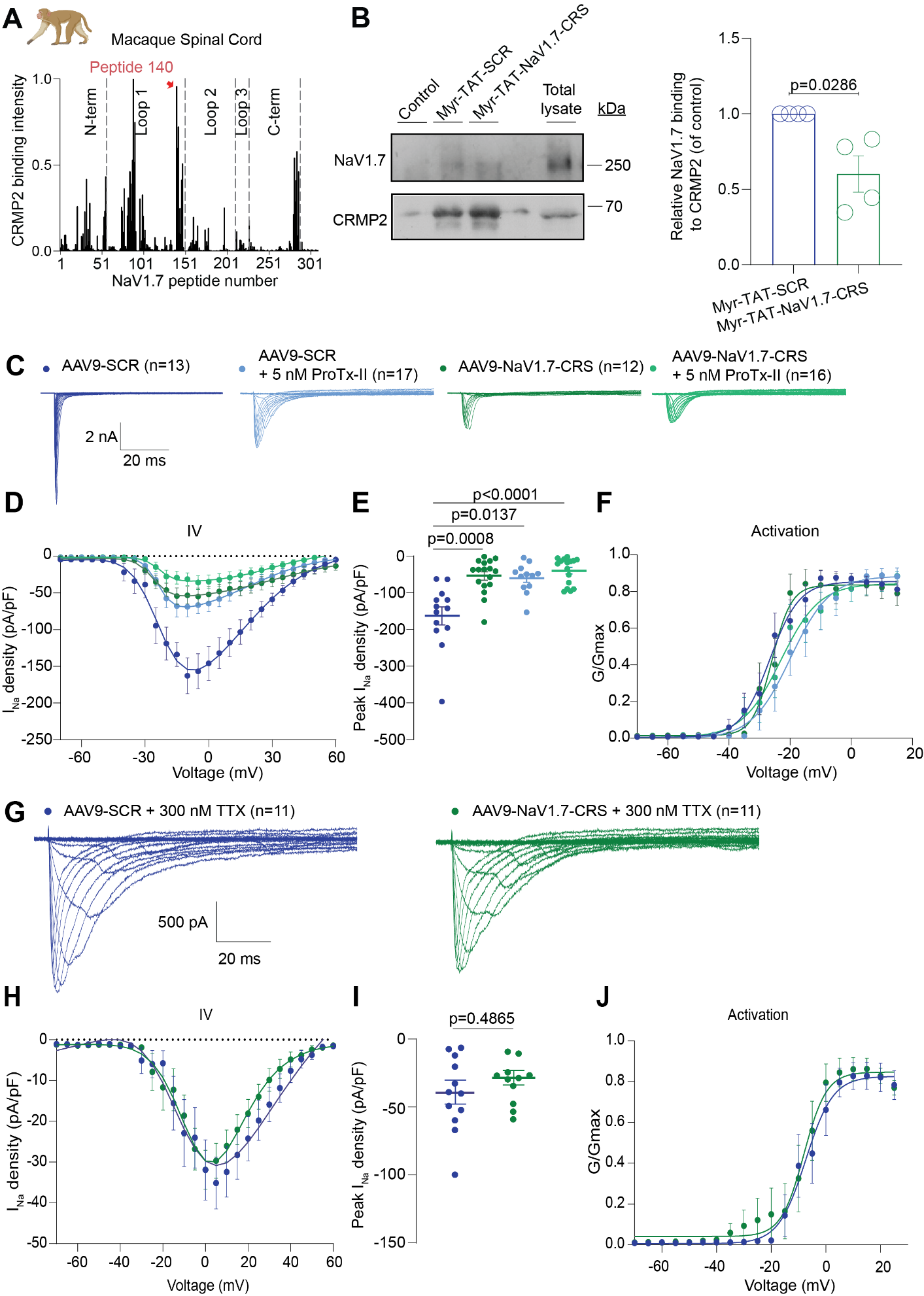
**Supplementary figure 10.** **Macaque DRG neurons transduced by AAV9-NaV1.7-CRS show reduced NaV1.7 currents and no effect on TTX-R currents.** (**A**) Fluorescent intensity of CRMP2 binding to the peptide array from macaque spinal cord lysate (n=2). The broadest peak with the highest CRMP2 binding, corresponding to peptide #140, is highlighted in red. (**B**) Representative immunoblots (left) and summary (right) of mean relative binding of NaV1.7 to CRMP2 in spinal cord lysates treated with the indicated peptides (n=4). Error bars show mean ± SEM; *p* values as indicated; Mann-Whitney test. (**C**) Representative current traces recorded from small-sized DRGs transduced with either AAV9-NaV1.7 or AAV9-SCR with co-application of 5 nM of the NaV1.7 specific inhibitor ProTx-II. (**D**) Boltzmann fits for current density–voltage curves. (**E**) Summary of peak current densities (pA/pF). (**F**) Boltzmann fits of voltage-dependent activation as shown. Half-maximal activation potential of activation and inactivation (*V_1/2_*) and slope values (*k*) for activation and inactivation are presented in **Table S4**. N=12-17 cells; error bars indicate mean ± SEM; *p* values as indicated; One-way ANOVA with Tukey’s post hoc test. (**G**) Representative current traces recorded from small-sized DRGs transduced with either AAV9-NaV1.7 or AAV9-SCR with co-application of 300 nM TTX. The remaining current fraction is due exclusively to TTX-R (NaV1.8 and NaV1.9) sodium channels. (**H**) Boltzmann fits for current density–voltage curves. (**I**) Summary of peak current densities (pA/pF). (**J**) Boltzmann fits of voltage-dependent activation as shown. Half-maximal activation potential of activation and inactivation (*V_1/2_*) and slope values (*k*) for activation and inactivation are presented in Table S3. N=11 cells; error bars indicate mean ± SEM; *p* values as indicated; One-way ANOVA with Tukey’s post hoc test.

**Table S1. Biophysical properties of Halo-NaV1.7(WT) channels and Halo-NaV1.7(X.X) mutants.**

|  | *Halo-NaV1.7(WT)* | *Halo-NaV1.7(1.1)* |
| --- | --- | --- |
| Activation |  |  |
| *V_1/2_* | -17.496 ± 1.198 (15) | -10.538 ± 1.063 (16)† |
| *k* | 7.471 ± 1.077 (15) | 8.204 ± 0.960 (16) |
| Inactivation |  |  |
| *V_1/2_* | -43.182 ± 2.203 (15) | -48.369 ± 1.667 (16) |
| *k* | -15.851 ± 2.138 (15) | -15.692 ± 1.662 (16) |
|  | *Halo-NaV1.7(WT)* | *Halo-NaV1.7(1.2)* |
| Activation |  |  |
| *V_1/2_* | -15.811 ± 0.638 (12) | -15.811 ± 1.277 (13) |
| *k* | 5.295 ± 0.446 (12) | 7.748 ± 1.156 (13) |
| Inactivation |  |  |
| *V_1/2_* | -47.307 ± 2.044 (12) | -45.667 ± 1.737 (13) |
| *k* | -15.826 ± 2.035 (12) | -15.612 ± 1.645 (13) |
|  | *Halo-NaV1.7(WT)* | *Halo-NaV1.7(1.3)* |
| Activation |  |  |
| *V_1/2_* | -15.196 ± 0.765 (15) | -17.324 ± 1.183 (16) |
| *k* | 6.165 ± 0.792 (15) | 6.491 ± 1.051 (16) |
| Inactivation |  |  |
| *V_1/2_* | -47.590 ± 3.319 (15) | -51.706 ± 3.260 (16) |
| *k* | -20.832± 3.698 (15) | -21.118 ± 3.776 (16) |
|  | *Halo-NaV1.7(WT)* | *Halo-NaV1.7(1.4)* |
| Activation |  |  |
| *V_1/2_* | -14.888 ± 1.221 (18) | -12.678 ± 0.561(20) |
| *k* | 5.888 ± 1.076 (18) | 4.161 ± 0.488 (20) |
| Inactivation |  |  |
| *V_1/2_* | -40.564 ± 1.410 (18) | -42.828 ± 1.432 (20) |
| *k* | -13.607 ± 1.304 (18) | -14.3.73 ± 1.355 (20) |
|  | *Halo-NaV1.7(WT)* | *Halo-NaV1.7(1.5)* |
| Activation |  |  |
| *V_1/2_* | -13.023 ± 0.791 (15) | -5.748 ± 0.507 (14)* |
| *k* | 5.504 ± 0.694(15) | 5.282 ± 0.440 (14) |
| Inactivation |  |  |
| *V_1/2_* | -43.750 ± 1.313(15) | -43.053 ± 1.473 (14) |
| *k* | -16.615 ± 1.296 (15) | -15.696 ± 1.424 (14) |
|  | *Halo-NaV1.7(WT)* | *Halo-NaV1.7(1.6)* |
| Activation |  |  |
| *V_1/2_* | -13.297 ± 0.916 (12) | -10.434 ± 0.862 (15) |
| *k* | 6.087 ± 0.808 (12) | 6.413 ± 0.759 (15) |
| Inactivation |  |  |
| *V_1/2_* | -33.958 ± 1.537 (10) | -34.189 ± 1.419 (15) |
| *k* | -12.141 ± 1.351 (10) | -13.077 ± 1.250 (15) |
|  | *Halo-NaV1.7(WT)* | *Halo-NaV1.7(1.8)* |
| Activation |  |  |
| *V_1/2_* | -10.540 ± 0.539 (15) | -11.535 ± 0.557 (17) |
| *k* | 3.992 ± 0.468 (15) | 4.189 ± 0.484 (17) |
| Inactivation |  |  |
| *V_1/2_* | -41.582 ± 1.901 (15) | -41.428 ± 1.612 (17) |
| *k* | -16.574 ± 1.838 (15) | -15.696 ± 1.538 (17) |
|  | *Halo-NaV1.7(WT)* | *Halo-NaV1.7(1.9)* |
| Activation |  |  |
| *V_1/2_* | -12.167 ± 0.852 (15) | -11.754 ± 0.743 (15) |
| *k* | 6.128 ± 0.750 (15) | 5.496 ± 0.651 (15) |
| Inactivation |  |  |
| *V_1/2_* | -53.446 ± 1.114 (15) | -53.160 ± 1.524 (15) |
| *k* | -12.789 ± 1.053 (15) | -13.964 ± 1.477 (15) |

Values are means ± SEM calculated from fits of the data from the indicated number of individual cells (in parentheses) to the Boltzmann equation; *V_1/2_* midpoint potential (mV) for voltage-dependent of activation or inactivation; *k*, slope factor. These values pertain to Fig. S1 and S2. Data were analyzed with Mann-Whitney test. *V_1/2_* † p<0.0001 Halo-NaV1.7(WT) vs Halo-NaV1.7(1.1); *p<0.0001 Halo-NaV1.7(WT) vs Halo-NaV1.7(1.5). DRG, dorsal root ganglia; DMSO, dimethylsulfoxide; ANOVA, analysis of variance.

**Table S2.** **Sodium channel gating properties following treatment with Myr-TAT-NaV1.7-CRS or Myr-TAT-SCR control peptide.**

|  | *Myr-TAT-SCR* |  | *Myr-TAT-NaV1.7-CRS* |  |
| --- | --- | --- | --- | --- |
| Activation |  |  |  |  |
| *V_1/2_* | -20.891 ± 0.491 (10) |  | -18.222 ± 1.057 (9) |  |
| *k* | 3.451 ± 0.427 (10) |  | 6.619 ± 0.945 (9) |  |
| Inactivation |  |  |  |  |
| *V_1/2_* | -49.661 ± 1.074 (14) |  | -48.744 ± 2.074 (13) |  |
| *k* | -11.625 ± 1.063 (14) |  | -15.436 ± 2.330 (13) |  |
|  | *Myr-TAT-SCR +*  *Vehicle* | *Myr-TAT-SCR +*  *5 nM ProTx-II* | *Myr-TAT-NaV1.7-CRS + Vehicle* | *Myr-TAT-NaV1.7-CRS +*  *5 nM ProTx-II* |
| Activation |  |  |  |  |
| *V_1/2_* | -21.796 ± 0.886 (12) | -11.790 ± 1.451 (11)† | -16.754 ± 0.688 (11) | -13.379 ± 1.549 (7) |
| *k* | 6.478 ± 0.792 (12) | 9.025 ± 1.292 (11) | 5.930 ± 0.608 (11) | 8.568 ± 1.398 (7) |
| Inactivation |  |  |  |  |
| *V_1/2_* | -55.752 ± 1.149 (11) | -56.185 ± 2.578 (11) | -61.788 ± 1.600 (10) | -65.578 ± 1.562 (6) |
| *k* | -10.245 ± 1.084 (11) | -13.761 ± 2.662 (11) | -10.594 ± 1.474 (10) | -10.875 ± 1.389 (6) |
|  | *Myr-TAT-SCR + siRNA-Control* | *Myr-TAT-SCR +*  *siRNA-CRMP2* | *Myr-TAT-NaV1.7-CRS + siRNA-Control* | *Myr-TAT-NaV1.7-CRS +*  *siRNA-CRMP2* |
| Activation |  |  |  |  |
| *V_1/2_* | -18.817 ± 1.031(8) | -16.842 ± 1.082 (10) | -14.997 ± 0.707(10) | -18.547 ± 1.0 (10) |
| *k* | 7.885 ± 0.945 (8) | 7.682 ± 0.983 (10) | 6.970 ± 0.632 (10) | 7.644 ± 0.912 (10) |
| Inactivation |  |  |  |  |
| *V_1/2_* | -58.378 ± 1.822 (10) | -64.102 ± 1.470 (7) | -62.284 ± 1.388 (10) | -62.143 ± 1.903 (11) |
| *k* | -13.012 ± 1.740 (10) | -11.331 ± 1.336 (7) | -11.090 ± 1.267 (10) | -13.542 ± 1.797(11) |
|  | *Myr-TAT-SCR + DMSO* | *Myr-TAT-SCR +*  *20 µM Pitstop2* | *Myr-TAT-NaV1.7-CRS + DMSO* | *Myr-TAT-NaV1.7-CRS +*  *20 µM Pitstop2* |
| Activation |  |  |  |  |
| *V_1/2_* | -18.909 ± 0.689 (14) | -14.236 ± 0.904 (7) | -17.872 ± 1.466 (11) | -18.871 ± 0.998 (13) |
| *k* | 4.940 ± 0.603 (14) | 6.915 ± 0.805 (7) | 8.347 ± 1.355 (11) | 7.629 ± 0.911 (13) |
| Inactivation |  |  |  |  |
| *V_1/2_* | -54.698 ± 1.188 (9) | -61.458 ± 1.217 (7) | -61.391 ± 0.780 (8) | -61.199 ± 0.997 (10) |
| *k* | -11.740 ± 1.116 (9) | -7.550 ± 1.072 (7) | -10.437 ± 0.707 (8) | -9.674 ± 0.897 (10) |

Values are means ± SEM calculated from fits of the data from the indicated number of individual cells (in parentheses) to the Boltzmann equation; *V_1/2_* midpoint potential (mV) for voltage-dependent of activation or inactivation; *k*, slope factor. These values pertain to Figure 3 and Figure 4. Only statistically significant differences are indicated within the table. Data were analyzed with Mann-Whitney test and one-way ANOVA with Tukey post hoc test.

† p<0.0001 comparing Myr-TAT-SCR + Vehicle vs Myr-TAT-SCR + 5 nM ProTx-II (one-way ANOVA with Tukey post hoc test). DRG, dorsal root ganglia; DMSO, dimethylsulfoxide; ANOVA, analysis of variance.

**Table S3.** **Biophysical properties of sodium currents following genetic delivery of interfering peptide in rat DRG neurons.**

|  | *AAV-SCR* |  | *AAV-NaV1.7-CRS* |  |
| --- | --- | --- | --- | --- |
| Activation |  |  |  |  |
| *V_1/2_* | -17.370 ± 0.491 (12) |  | -17.945 ± 1.057 (12) |  |
| *k* | 8.232 ± 0.427 (12) |  | 8.028 ± 0.945 (12) |  |
| Inactivation |  |  |  |  |
| *V_1/2_* | -50.230± 1.074 (12) |  | -49.320 ± 2.074 (12) |  |
| *k* | -11.890 ± 1.063 (12) |  | -13.061 ± 2.330 (12) |  |
|  | *AAV-SCR +*  *Vehicle* | *AAV-SCR +*  *5 nM ProTx-II* | *AAV-NaV1.7-CRS + Vehicle* | *AAV-NaV1.7-CRS +*  *5 nM ProTx-II* |
| Activation |  |  |  |  |
| *V_1/2_* | -24.449 ± 0.620 (15) | -12.156 ± 0.922 (12)‡ | -26.747 ± 0.858 (13) | -24.001 ± 1.311 (15) |
| *k* | 5.797 ± 0.546 (15) | 7.247 ± 0.823 (12) | 5.349 ± 0.752 (13) | 8.170 ± 1.207 (15) |
| Inactivation |  |  |  |  |
| *V_1/2_* | -52.147 ± 1.614 (15) | -44.326 ± 2.692 (12)† | -58.741 ± 1.605 (13) | -52.516 ± 2.214 15) |
| *k* | -18.065 ± 1.728 (15) | -18.956 ± 2.78 (12) | -15.255 ± 1.598 (13) | -18.334 ± 2.389 (15) |
|  | *AAV-SCR + DMSO* | *AAV-SCR +*  *20 µM Pitstop2* | *AAV-NaV1.7-CRS + DMSO* | *AAV-NaV1.7-CRS +*  *20 µM Pitstop2* |
| Activation |  |  |  |  |
| *V_1/2_* | -21.946 ± 1.720 (12) | -25.782 ± 1.874 (15) | -23.054 ± 1.262 (15) | -17.621 ± 2.441 (9) |
| *k* | 8.521 ± 1.598 (12) | 9.756 ± 1.797 (15) | 8.063 ± 1.159 (15) | 11.247 ± 2.432 (9) |
| Inactivation |  |  |  |  |
| *V_1/2_* | -49.001 ± 1.848 (12) | -47.284 ± 1.497 (15)* | -55.146 ± 1.664 (15) | -48.747 ± 1.592 (9) |
| *k* | -15.754 ± 1.850 (12) | -14.758 ± 1.457 (15) | -14.475 ± 1.634 (15) | -13.477 ± 1.518 (9) |

Values are means ± SEM calculated from fits of the data from the indicated number of individual cells (in parentheses) to the Boltzmann equation; *V_1/2_* midpoint potential (mV) for voltage-dependent of activation or inactivation; *k*, slope factor. These values pertain to biophysical properties of recordings from Fig. 7. Only statistically significant differences are indicated within the table. Data were analyzed with Mann-Whitney test and one-way ANOVA with Tukey post hoc test.

† p= 0.0493 for *V_1/2_* inactivation of AAV-SCR + ProTx-II 5nM vs AAV-SCR Vehicle. p<0.0363 AAV-SCR + ProTx-II vs AAV-NaV1.7-CRS+ ProTx-II, p<0.0001 AAV-SCR + ProTx-II5 nM vs. AAV-NaV1.7-CRS+ ProTx-II 5 nM (one-way ANOVA with Tukey post hoc test). ‡ p<0.0001 comparing AAV-SCR + Water vs AAV-SCR + 5 nM ProTx-II, and AAV-NaV1.7-CRS+ Vehicle, and AAV-NaV1.7-CRS+ ProTx-II (one-way ANOVA with Tukey post hoc test). * p=0.0042 for *V_1/2_* inactivation AAV-SCR + Pitstop 20 µM vs. AAV-NaV1.7-CRS+ Vehicle. DRG, dorsal root ganglia; DMSO, dimethylsulfoxide; ANOVA, analysis of variance.

**Table S4.** **Biophysical properties of sodium currents following genetic delivery of interfering peptide in macaque DRG neurons.**

|  | *AAV9-SCR + TTX* |  | *AAV9-NaV1.7-CRS+ TTX* |  |
| --- | --- | --- | --- | --- |
| Activation |  |  |  |  |
| *V_1/2_* | -7.361 ± 1.232 (11) |  | -8.037 ± 1.238 (11) |  |
| *k* | 4.992 ± 1.072 (11) |  | 4.151 ± 1.074 (11) |  |
|  | *AAV9-SCR +*  *Vehicle* | *AAV9-SCR +*  *5 nM ProTx-II* | *AAV9-NaV1.7-CRS+ Vehicle* | *AAV9-NaV1.7-CRS+*  *5 nM ProTx-II* |
| Activation |  |  |  |  |
| *V_1/2_* | -26.624 ± 1.017 (13) | -19.767 ± 1.500 (13)† | -25.920 ± 0.890 (17) | -23.060 ± 1.480 (14) |
| *k* | 4.824 ± 0.892 (13) | 6.424 ± 1.339 (13) | 3.177 ± 0.771 (17) | 6.320 ± 1.327 (14) |

Values are means ± SEM calculated from fits of the data from the indicated number of individual cells (in parentheses) to the Boltzmann equation; *V_1/2_* midpoint potential (mV) for voltage-dependent of activation or inactivation; *k*, slope factor. These values pertain to biophysical properties of recordings from Fig. 7. Only statistically significant differences are indicated within the table. Data were analyzed with Mann-Whitney test and one-way ANOVA with Tukey post hoc test.

† p= 0.0037 for *V_1/2_* activation of AAV9-CMV-SCR vs. AAV9-CMV-SCR + ProTx-II. p=0.0175 AAV9-CMV-SCR + ProTx-II vs. AAV9-CMV-NaV1.7 (one-way ANOVA with Tukey post hoc test). DRG, dorsal root ganglia; DMSO, dimethylsulfoxide; ANOVA, analysis of variance.

**Table S5. Details of statistical comparisons.**

| **Figure panel** | **Assay** | **Statistical Test; findings** | **Post-hoc analysis (adjusted p-values)** | **Number of subjects** | **Number of subjects excluded (ROUT test)** |
| --- | --- | --- | --- | --- | --- |
| Fig 1G | Whole cell patch clamp electrophysiology – Peak sodium currents | One Way ANOVA | Holm-Sidak multiple comparison post-hoc test:  Halo-NaV1.7(WT) + TTX vs. Halo-NaV1.7(WT) + TTX + ProTx-II  p=0.0017  Halo-NaV1.7(WT) + TTX vs. Halo-NaV1.7(1.3) + TTX  p=0.0003  Halo-NaV1.7(WT) + TTX vs. Halo-NaV1.7(1.3) + TTX + ProTx-II  p=0.0006 | Halo-NaV1.7(WT) + TTX (n=16)  Halo-NaV1.7(WT) + TTX + ProTx-II  (n=11)  Halo-NaV1.7(1.3) + TTX  (n=15)  Halo-NaV1.7(1.3) + TTX + ProTx-II  (n=11) | Halo-NaV1.7(WT) + TTX excluded n=1 |
| Fig 1H | Whole cell patch clamp electrophysiology – Peak TTX-R currents | Kruskal-Wallis test | Dunn’s multiple comparison post-hoc test:  Halo-NaV1.7(WT) + TTX vs. Halo-NaV1.7(WT) + TTX + ProTx-II  p >0.9999  Halo-NaV1.7(WT) + TTX vs. Halo-NaV1.7(1.3) + TTX  p>0.9999  Halo-NaV1.7(WT) + TTX vs. Halo-NaV1.7(1.3) + TTX + ProTx-II  p>0.9999 | Halo-NaV1.7(WT) + TTX (n=16)  Halo-NaV1.7(WT) + TTX + ProTx-II  (n=11)  Halo-NaV1.7(1.3) + TTX  (n=15)  Halo-NaV1.7(1.3) + TTX + ProTx-II  (n=11) | Halo-NaV1.7(WT) + TTX excluded n=1 |
| Fig 2C | CRMP2 immunoprecipitation followed by Nav1.7 western blot | Mann Whitney | DMSO vs. Myr-TAT-SCR p>0.9999; DMSO vs. Myr-TAT-NaV1.7-CRS p=0.0267 | DMSO (n=5)  Myr-TAT-SCR (n=5)  Myr-TAT-NaV1.7-CRS (n=5) |  |
| Fig 2F | Whole cell patch clamp electrophysiology – peak current density | Welch’s t-test | Myr-TAT-NaV1.7-CRS vs.  Myr-TAT-SCR  p=0.0396 | Myr-TAT-NaV1.7-CRS (N=15); Myr-TAT-SCR (N=13) |  |
| Fig 2G | Whole cell patch clamp electrophysiology – Peak TTX-R current | Mann-Whitney test | Myr-TAT-NaV1.7-CRS vs.  Myr-TAT-SCR  p=0.8460 | Myr-TAT-NaV1.7-CRS  (N=16)  Myr-TAT-SCR (N=13) |  |
| Fig 2J | Whole cell patch clamp electrophysiology – peak current density | One-way ANOVA | Tukey’s multiple comparison post-hoc test:  Myr-TAT-SCR + Water vs.  Myr-TAT-SCR + 5 nM ProTx-II  p=0.0130  Myr-TAT-SCR + Water vs.  Myr-TAT-NaV1.7-CRS + Water  p=0.0470  Myr-TAT-SCR + Water vs.  Myr-TAT-NaV1.7-CRS + 5 nM ProTx-II  p=0.0012  Myr-TAT-NaV1.7-CRS + Water vs.  Myr-TAT-NaV1.7-CRS + 5 nM ProTx-II  p=0.8197 | Myr-TAT-SCR + Water (N=14)  Myr-TAT-SCR + 5 nM ProTx-II (N=13)  Myr-TAT-NaV1.7-CRS + Water (N=12)  Myr-TAT-NaV1.7-CRS + 5 nM ProTx-II (N=8) |  |
| Fig 2M | Whole cell patch clamp electrophysiology – peak current density | One-way ANOVA | Tukey’s multiple comparison post-hoc test:  Myr-TAT-SCR + Ctrl siRNA vs. Myr-TAT-SCR + CRMP2 siRNA  p=0.0416  Myr-TAT-SCR + Ctrl siRNA vs. Myr-TAT-NaV1.7-CRS + Ctrl siRNA  p=0.0573  Myr-TAT-SCR + Ctrl siRNA vs. Myr-TAT-NaV1.7-CRS + CRMP2 siRNA  p=0.0499  Myr-TAT-NaV1.7-CRS + Ctrl siRNA vs. Myr-TAT-NaV1.7-CRS + CRMP2 siRNA p>0.9999 | Myr-TAT-SCR + Ctrl siRNA (n=13)  Myr-TAT-SCR + CRMP2 siRNA (n=12)  Myr-TAT-NaV1.7-CRS + Ctrl siRNA (n=12)  Myr-TAT-NaV1.7-CRS + CRMP2 siRNA  (n=13) |  |
| Fig 2O | Total protein western blot | Kruskal-Wallis test | DMSO vs. Myr-TAT-SCR  p=0.4620  DMSO vs. Myr-TAT-NaV1.7-CRS  p>0.9999 | DMSO (n=4)  Myr-TAT-SCR (n=4)  Myr-TAT-NaV1.7-CRS (n=4) |  |
| Fig 2Q | Cell surface biotinylation | One-way ANOVA | Kruskal-Wallis test:  DMSO vs. Myr-TAT-SCR  p=0.8734  DMSO vs. Myr-TAT-NaV1.7-CRS  p=0.0178 | DMSO (n=5)  Myr-TAT-SCR (n=5)  Myr-TAT-NaV1.7-CRS (n=5) |  |
| Fig 2T | Whole cell patch clamp electrophysiology – peak current density | One-way ANOVA  p=0.0022 | Kurskal-Wallis multiple comparison post-hoc test:  Myr-TAT-SCR peptide + 0.1% DMSO  vs.  Myr-TAT-SCR peptide + 20 µM Pitstop2  p>0.9999  Myr-TAT-SCR peptide + 0.1% DMSO  vs.  Myr-TAT-NaV1.7-CRS peptide + 0.1% DMSO  p=0.0033  Myr-TAT-SCR peptide + 20uM Pitstop2  vs.  Myr-TAT-NaV1.7-CRS peptide + 20 µM Pitstop2  p>0.9999  Myr-TAT-NaV1.7-CRS peptide + 0.1% DMSO  vs.  Myr-TAT-NaV1.7-CRS peptide + 20 µM Pitstop2  p=0.0109 | Myr-TAT-SCR peptide + 0.1% DMSO (n=15)  Myr-TAT-SCR peptide + 20uM Pitstop2 (n=7)  Myr-TAT-NaV1.7-CRS peptide + 0.1% DMSO (n=12)  Myr-TAT-NaV1.7-CRS peptide + 20uM Pitstop2 (n=14) |  |
| Fig 3B | Whole cell patch clamp electrophysiology – evoked action potentials | Multiple Mann-Whitney tests | Myr-TAT-SCR vs. Myr-TAT-NaV1.7-CRS  0 pA, p>0.9999  10 pA, p=0.2121  20 pA, p=0.0644  30 pA, p=0.0063  40 pA, p=0.0063  50 pA, p=0.0076  60 pA, p=0.0189  70 pA, p=0.0013  80 pA, p=0.0088  90 pA, p=0.0063  100 pA, p=0.0088  110 pA, p=0.0088  120 pA, p=0.0101 | Myr-TAT-SCR (n=5)  Myr-TAT-NaV1.7-CRS (n=7) |  |
| Fig 3C | Whole cell patch clamp electrophysiology – resting membrane potential | Welch’s t-test | Myr-TAT-SCR vs. Myr-TAT-NaV1.7-CRS  p=0.7603 | Myr-TAT-SCR (n=5)  Myr-TAT-NaV1.7-CRS (n=7) |  |
| Fig 3E | Whole cell patch clamp electrophysiology – rheobase | Welch’s t-test | Myr-TAT-SCR vs. Myr-TAT-NaV1.7-CRS  p=0.0308 | Myr-TAT-SCR (n=5)  Myr-TAT-NaV1.7-CRS (n=7) |  |
| Fig 3H | Spinal cord CGRP release assay | Two-Way ANOVA | Sidak’s multiple comparisons test  Fraction 1  Control vs. Myr-TAT-SCR  p=0.8968  Control vs. Myr-TAT-NaV1.7-CRS p=0.1324  Myr-TAT-SCR vs Myr-TAT-NaV1.7-CRS p=0.4010  Fraction 2  Control vs. Myr-TAT-SCR  p=0.2430  Control vs. Myr-TAT-NaV1.7-CRS p=0.1495  Myr-TAT-SCR vs. Myr-TAT-NaV1.7-CRS p=0.8924  Fraction 3  Control vs. Myr-TAT-SCR  p=0.9219  Control vs. Myr-TAT-NaV1.7-CRS p=0.0026  Myr-TAT-SCR vs. Myr-TAT-NaV1.7-CRS p=0.0012  Fraction 4  Control vs. Myr-TAT-SCR  p=0.2418  Control vs. Myr-TAT-NaV1.7-CRS p<0.0001  Myr-TAT-SCR vs. Myr-TAT-NaV1.7-CRS p<0.0001  Fraction 5  Control vs. Myr-TAT-SCR  p=0.5405  Control vs. Myr-TAT-NaV1.7-CRS p=0.0191  Myr-TAT-SCR vs. Myr-TAT-NaV1.7-CRS p=0.4724  Fraction 6  Control vs. Myr-TAT-SCR  p=0.9613  Control vs. Myr-TAT-NaV1.7-CRS p=0.7229  Myr-TAT-SCR vs. Myr-TAT-NaV1.7-CRS p=0.9202 | DMSO (n=4)  Myr-TAT-SCR (n=4)  Myr-TAT-NaV1.7-CRS (n=4) |  |
| Fig 4A | CRMP2 binding to peptide 141 | Mann-Whitney test | WT vs. CRMP2^K374A/K374A^  p<0.0001 | WT (n=4)  CRMP2^K374A/K374A^ (n=4) |  |
| Fig 4B | SNI female rats – paw withdrawal threshold | Multiple Mann-Whitney tests | Myr-TAT-SCR female vs. Myr-TAT-NaV1.7-CRS female (Pre-SNI, p>0.9999)  Myr-TAT-SCR female vs. Myr-TAT-NaV1.7-CRS female (t=0, p=0.9675)  Myr-TAT-SCR female vs. Myr-TAT-NaV1.7-CRS female (t=0.5, p>0.9999)  Myr-TAT-SCR female vs. Myr-TAT-NaV1.7-CRS female (t=1, p=0.8918)  Myr-TAT-SCR female vs. Myr-TAT-NaV1.7-CRS female (t=2, p=0.0173)  Myr-TAT-SCR female vs. Myr-TAT-NaV1.7-CRS female (t=3, p=0.0022)  Myr-TAT-SCR female vs. Myr-TAT-NaV1.7-CRS female (t=4, p=0.0152)  Myr-TAT-SCR female vs. Myr-TAT-NaV1.7-CRS female (t=5, p=0.0649) | Myr-TAT-SCR female (n=6)  Myr-TAT-NaV1.7-CRS female (n=6) |  |
|  | SNI male rats – paw withdrawal threshold |  | Myr-TAT-SCR male vs. Myr-TAT-NaV1.7-CRS male (Pre-SNI, p>0.9999)  Myr-TAT-SCR male vs. Myr-TAT-NaV1.7-CRS male (t=0, p=0.3095)  Myr-TAT-SCR male vs. Myr-TAT-NaV1.7-CRS male (t=0.5, p=0.4643)  Myr-TAT-SCR male vs. Myr-TAT-NaV1.7-CRS male (t=1, p=0.0833)  Myr-TAT-SCR male vs. Myr-TAT-NaV1.7-CRS male (t=2, p=0.0119)  Myr-TAT-SCR male vs. Myr-TAT-NaV1.7-CRS male (t=3, p=0.0119)  Myr-TAT-SCR male vs. Myr-TAT-NaV1.7-CRS male (t=4, p=0.0119)  Myr-TAT-SCR male vs. Myr-TAT-NaV1.7-CRS male (t=5, p=0.0119) | Myr-TAT-SCR male (n=6)  Myr-TAT-NaV1.7-CRS male (n=6) |  |
| Fig 4C | Paw withdrawal threshold rats – area under the curve | Mann-Whitney test | Myr-TAT-SCR female vs. Myr-TAT-NaV1.7-CRS female  p=0.0087  Myr-TAT-SCR male vs. Myr-TAT-NaV1.7-CRS male  p=0.0152 | Myr-TAT-SCR female (n=6)  Myr-TAT-NaV1.7-CRS female (n=6)  Myr-TAT-SCR male (n=6)  Myr-TAT-NaV1.7-CRS male (n=6) |  |
| Fig 4D | Mouse thermal nociception – hot plate | Kruskal-Wallis test | Myr-TAT-SCR vs. Myr-TAT-NaV1.7-CRS (females) p=0.1177  Myr-TAT-SCR vs. Myr-TAT-NaV1.7-CRS (males) p=0.5540 | Female Myr-TAT-SCR (n=10)  Female Myr-TAT-NaV1.7-CRS (n=10)  Male Myr-TAT-SCR (n=10)  Male Myr-TAT-NaV1.7-CRS (n=10) |  |
| Fig 4E | Mouse thermal nociception – tail flick | Kruskal-Wallis test | Myr-TAT-SCR vs. Myr-TAT-NaV1.7-CRS (females) p>0.9999  Myr-TAT-SCR vs. Myr-TAT-NaV1.7- CRS (males) p>0.9999 | Female Myr-TAT-SCR (n=10)  Female Myr-TAT-NaV1.7-CRS (n=10)  Male Myr-TAT-SCR (n=10)  Male Myr-TAT-NaV1.7-CRS (n=10) |  |
| Fig 4F | Motor coordination in rats -rotarod | Multiple Mann-Whitney tests | Males  Myr-TAT-SCR vs. Myr-TAT-NaV1.7-CRS (baseline) p=0.4009  Myr-TAT-SCR vs. Myr-TAT-NaV1.7-CRS (60 min post injection) p=0.8298  Myr-TAT-SCR vs. Myr-TAT-NaV1.7-CRS (120 min post injection) p=0.3176  Myr-TAT-SCR vs. Myr-TAT-NaV1.7-CRS (180 min post injection) p=0.7103  Myr-TAT-SCR vs. Myr-TAT-NaV1.7-CRS (240 min post injection) p=0.9015  Myr-TAT-SCR vs. Myr-TAT-NaV1.7-CRS (300 min post injection) p=0.6859 | Myr-TAT-SCR (n=7)  Myr-TAT-NaV1.7-CRS (n=7) |  |
| Fig 5C | Whole cell patch clamp electrophysiology – peak current density | Welch’s t-test | Myr-TAT-SCR vs. Myr-TAT-NaV1.7-CRS p=0.0002 | Myr-TAT-SCR (n=12)  Myr-TAT-NaV1.7-CRS (n=12) |  |
| Fig 5D | Whole cell patch clamp electrophysiology – TTX-R current density | Mann-Whitney test | Myr-TAT-SCR vs. Myr-TAT-NaV1.7-CRS  p=0.6194 | Myr-TAT-SCR (n=12)  Myr-TAT-NaV1.7-CRS (n=12) |  |
| Fig 5G | Whole cell patch clamp electrophysiology – peak current density | One-way ANOVA | pAAV-SCR +Vehicle vs.  pAAV-SCR + ProTx  p=0.0024  pAAV-SCR +Vehicle vs. pAAV-NaV1.7-CRS + vehicle  p=0.0213  pAAV-SCR +Vehicle vs. pAAV-NaV1.7-CRS + ProTx  p=0.0003  pAAV-NaV1.7-CRS + Vehicle vs. pAAV-NaV1.7-CRS + ProTx  p>0.9999 | pAAV-SCR +Vehicle (n=15)  pAAV-SCR + Protox (n=12)  pAAV-NaV1.7-CRS + Vehicle (n=13)  pAAV-NaV1.7-CRS + Protox (n=15) |  |
| Fig 5J | Whole cell patch clamp electrophysiology – peak current density | One-way ANOVA | pAAV-SCR +Vehicle vs. pAAV-SCR + Pitstop2  p>0.9999  pAAV-SCR +Vehicle vs. pAAV-NaV1.7-CRS + Vehicle  p>0.0018  pAAV-SCR +Vehicle vs. pAAV-NaV1.7-CRS + Pitstop2  p>0.9999  pAAV-NaV1.7-CRS + Vehicle v.s pAAV-NaV1.7-CRS + Pitstop2  p=0.0115 | pAAV-SCR +Vehicle (n=12)  pAAV-SCR + Pitstop2 (n=15)  pAAV-NaV1.7-CRS + Vehicle (n=15)  pAAV-NaV1.7-CRS + Pitstop2 (n=9) |  |
| Fig 5L | Slice patch clamp electrophysiology – Frequency of sEPSC | Mann-Whitney test | pAAV-SCR vs. pAAV-NaV1.7-CRS  p=0.0305 | pAAV-SCR (n=12)  pAAV-NaV1.7-CRS (n=11) |  |
| Fig 5L | Slice patch clamp electrophysiology – Amplitud of sEPSC | Mann-Whitney test | pAAV-SCR vs. pAAV-NaV1.7-CRS  p=0.0357 | pAAV-SCR (n=11)  pAAV-NaV1.7-CRS (n=10) |  |
| Fig 6D | Males SNI reversal – paw withdrawal thresholds | Two-way anova | AAV-CMV-eGFP-NaV1.7-CRS vs. AAV-CMV-eGFP-SCR (Pre-SNI, p>0.9999)  AAV-CMV-eGFP-NaV1.7-CRS vs. AAV-CMV-eGFP-SCR (t=0d, p>0.9999)  AAV-CMV-eGFP-NaV1.7-CRS vs. AAV-CMV-eGFP-SCR (t=4d, p=0.0173)  AAV-CMV-eGFP-NaV1.7-CRS vs. AAV-CMV-eGFP-SCR (t=7d, p=0.0087)  AAV-CMV-eGFP-NaV1.7-CRS vs. AAV-CMV-eGFP-SCR (t=11d, p=0.0152)  AAV-CMV-eGFP-NaV1.7-CRS vs. AAV-CMV-eGFP-SCR (t=14d, p=0.0087)  AAV-CMV-eGFP-NaV1.7-CRS vs. AAV-CMV-eGFP-SCR (t=20d, p=0.0238)  AAV-CMV-eGFP-NaV1.7-CRS vs. AAV-CMV-eGFP-SCR (t=24d, p=0.0866)  AAV-CMV-eGFP-NaV1.7-CRS vs. AAV-CMV-eGFP-SCR (t=31d, p=0.1212) | AAV-CMV-eGFP-NaV1.7-CRS (n=5)  AAV-CMV-eGFP-SCR (n=6) |  |
| Fig 6E | Males SNI reversal – area under the curve | Mann-Whitney test | AAV-CMV-eGFP-NaV1.7-CRS vs AAV-CMV-eGFP-SCR (p=0.0043) | AAV-CMV-eGFP-NaV1.7-CRS (n=5)  AAV-CMV-eGFP-SCR (n=6) |  |
| Fig 6F | Males SNI- Reversal  Locomotor activity | Mann-Whitney test | AAV9-CMV-eGFP-SCR vs. AAV9-CMV-eGFP-NaV1.7  p=0.9307 | AAV9-CMV-eGFP-SCR  (n=6)  AAV9-CMV-eGFP-NaV1.7  (n=5) |  |
| Fig 6G | Males SNI- Reversal  Open field test | Multiple Mann-Whitney tests | Time in Periphery  AAV9-CMV-eGFP-SCR vs. AAV9-CMV-eGFP-NaV1.7  p=0.2786  Time in Center  AAV9-CMV-eGFP-SCR vs. AAV9-CMV-eGFP-NaV1.7  p=0.2786 | AAV9-CMV-eGFP-SCR  (n=6)  AAV9-CMV-eGFP-NaV1.7  (n=5) |  |
| Fig 6H | Females SNI reversal – paw withdrawal thresholds | Two-way anova | AAV-CMV-eGFP-NaV1.7-CRS vs. AAV-CMV-eGFP-SCR (Pre-SNI, p>0.9999)  AAV-CMV-eGFP-NaV1.7-CRS vs. AAV-CMV-eGFP-SCR (t=7d, p=0.3770)  AAV-CMV-eGFP-NaV1.7-CRS vs. AAV-CMV-eGFP-SCR (t=11d, p=0.0275)  AAV-CMV-eGFP-NaV1.7-CRS vs. AAV-CMV-eGFP-SCR (t=14d, p=0.021)  AAV-CMV-eGFP-NaV1.7-CRS vs. AAV-CMV-eGFP-SCR (t=18d, p<0.0001)  AAV-CMV-eGFP-NaV1.7-CRS vs. AAV-CMV-eGFP-SCR (t=27d, p<0.0001)  AAV-CMV-eGFP-NaV1.7-CRS vs. AAV-CMV-eGFP-SCR (t=38d, p=0.004) | AAV-CMV-eGFP-SCR (n=8)  AAV-CMV-eGFP-NaV1.7-CRS (n=9) |  |
| Fig 6I | Females SNI reversal – area under the curve | Mann-Whitney test | AAV-CMV-eGFP-NaV1.7-CRS vs. AAV-CMV-eGFP-SCR (p<0.0001) | AAV-CMV-eGFP-SCR (n=8)  AAV-CMV-eGFP-NaV1.7-CRS (n=9) |  |
| Fig 6J | Females SNI reversal - Locomotor activity | Mann-Whitney test | AAV9-CMV-eGFP-SCR vs. AAV9-CMV-eGFP-NaV1.7  p=0.1520 | AAV9-CMV-eGFP-SCR  (n=7)  AAV9-CMV-eGFP-NaV1.7  (n=8) |  |
| Fig 6K | Females SNI- reversal  Open field test | Multiple Mann-Whitney tests | Time in Periphery  AAV9-CMV-eGFP-SCR vs. AAV9-CMV-eGFP-NaV1.7  p=0.8665  Time in Center  AAV9-CMV-eGFP-SCR vs. AAV9-CMV-eGFP-NaV1.7  p=0.8665 |  |  |
| Fig 6L | Males SNI prevention – paw withdrawal thresholds | Two-way anova | AAV-CMV-eGFP-NaV1.7-CRS vs. AAV-CMV-eGFP-SCR (BL1, p>0.9999)  AAV-CMV-eGFP-NaV1.7-CRS vs. AAV-CMV-eGFP-SCR (BL2, p>0.9999)  AAV-CMV-eGFP-NaV1.7-CRS vs. AAV-CMV-eGFP-SCR (t=6d, p=0.0424)  AAV-CMV-eGFP-NaV1.7-CRS vs. AAV-CMV-eGFP-SCR (t=8d, p=0.0002)  AAV-CMV-eGFP-NaV1.7-CRS vs. AAV-CMV-eGFP-SCR (t=12d, p=0.0005)  AAV-CMV-eGFP-NaV1.7-CRS vs. AAV-CMV-eGFP-SCR (t=19d, p=0.0005)  AAV-CMV-eGFP-NaV1.7-CRS vs. AAV-CMV-eGFP-SCR (t=21d, p=0.0015)  AAV-CMV-eGFP-NaV1.7-CRS vs. AAV-CMV-eGFP-SCR (t=54d, p=0.0038) | AAV-CMV-eGFP-SCR (n=7)  AAV-CMV-eGFP-NaV1.7-CRS (n=9) |  |
| Fig 6M | Males SNI prevention – area under the curve | Mann-Whitney test | AAV-CMV-eGFP-NaV1.7-CRS vs. AAV-CMV-eGFP-SCR (p=0.0002) | AAV-CMV-eGFP-SCR (n=7)  AAV-CMV-eGFP-NaV1.7-CRS (n=9) |  |
| Fig 6N | Males SNI prevention - Locomotor activity | Mann-Whitney test | AAV9-CMV-eGFP-SCR vs. AAV9-CMV-eGFP-NaV1.7  p=0.7577 | AAV-CMV-eGFP-SCR (n=7)  AAV-CMV-eGFP-NaV1.7-CRS (n=9) |  |
| Fig 6O | Males SNI- Prevention  Open field test | Multiple Mann-Whitney tests | Time in Periphery  AAV9-CMV-eGFP-SCR vs. AAV9-CMV-eGFP-NaV1.7  P=0.1141  Time in Center  AAV9-CMV-eGFP-SCR vs. AAV9-CMV-eGFP-NaV1.7  P=0.1141 | AAV-CMV-eGFP-SCR (n=7)  AAV-CMV-eGFP-NaV1.7-CRS (n=9) |  |
| Fig 6P | Females SNI prevention – paw withdrawal thresholds | Two-way anova | AAV-CMV-eGFP-NaV1.7-CRS vs. AAV-CMV-eGFP-SCR (BL1, p>0.9999)  AAV-CMV-eGFP-NaV1.7-CRS vs. AAV-CMV-eGFP-SCR (BL2, p=0.2058)  AAV-CMV-eGFP-NaV1.7-CRS vs. AAV-CMV-eGFP-SCR (t=8d, p=0.0024)  AAV-CMV-eGFP-NaV1.7-CRS vs. AAV-CMV-eGFP-SCR (t=12d, p=0.0005)  AAV-CMV-eGFP-NaV1.7-CRS vs. AAV-CMV-eGFP-SCR (t=16d, p=0.0256)  AAV-CMV-eGFP-NaV1.7-CRS vs. AAV-CMV-eGFP-SCR (t=20d, p=0.0069)  AAV-CMV-eGFP-NaV1.7-CRS vs. AAV-CMV-eGFP-SCR (t=54d, p=0.0107) | AAV-CMV-eGFP-SCR (n=9)  AAV-CMV-eGFP-NaV1.7-CRS (n=8) |  |
| Fig 6Q | Females SNI prevention – area under the curve | Mann-Whitney test | AAV-CMV-eGFP-NaV1.7-CRS vs AAV-CMV-eGFP-SCR (p<0.0001) | AAV-CMV-eGFP-SCR (n=9)  AAV-CMV-eGFP-NaV1.7-CRS (n=8) |  |
| Fig 6R | Females SNI prevention - Locomotor activity | Mann-Whitney test | AAV9-CMV-eGFP-SCR vs. AAV9-CMV-eGFP-NaV1.7  p=0.3704 | AAV-CMV-eGFP-SCR (n=9)  AAV-CMV-eGFP-NaV1.7-CRS (n=8) |  |
| Fig 6S | Females SNI- prevention  Open field test | Multiple Mann-Whitney tests | Time in Periphery  AAV9-CMV-eGFP-SCR vs. AAV9-CMV-eGFP-NaV1.7  p=0.5414  Time in Center  AAV9-CMV-eGFP-SCR vs. AAV9-CMV-eGFP-NaV1.7 p=0.5414 | AAV-CMV-eGFP-SCR (n=9)  AAV-CMV-eGFP-NaV1.7-CRS (n=8) |  |
